# Bayesian inference of multi-point macromolecular architecture mixtures at nanometre resolution

**DOI:** 10.1101/2021.08.22.457266

**Authors:** Peter A Embacher, Tsvetelina E Germanova, Emanuele Roscioli, Andrew D McAinsh, Nigel J Burroughs

## Abstract

Gaussian spot fitting methods have significantly extended the spatial range where fluorescent microscopy can be used, with recent techniques approaching nanometre (nm) resolutions. However, small inter-fluorophore distances are systematically over-estimated for typical molecular scales (≲ 50nm). This bias can be corrected computationally, but current algorithms are limited to correcting distances between pairs of fluorophores. Here we present a flexible Bayesian computational approach that infers the distances and angles between multiple markers and has several advantages over these previous methods. Specifically it improves confidence intervals for small lengths, estimates measurement errors of each fluorescent marker individually and infers the correlations between polygon lengths. The latter is essential for determining the full multi-fluorophore 3D architecture. We further developed the algorithm to infer the mixture composition of a heterogeneous population of multiple polygon states. We use our algorithm to analyse the 3D architecture of the human kinetochore, a macro-molecular complex that is essential for high fidelity cell division. We examine the conformational change induced by microtubule attachment using triple fluorophore marked data and demonstrate for the first time that in metaphase kinetochore conformation is heterogeneous.

## Introduction

The classical Rayleigh criterion limits the resolution of light microscopy to about 200nm for typical wavelengths and numerical apertures. However, this limit can in principle be pushed arbitrarily close to zero by fitting the point spread function (PSF) to diffraction limited objects [11], [21]. In this case the localisation accuracy is primarily limited by the finite signal to noise (S/N) ratio, a consequence of the finite photon count [19], [29]. By using multiple fluorophores to label diffraction limited objects, inter-object distances can thus be measured, achieving a localisation accuracy of typically tens of nm with standard fluorescence microscopes and markers.

Pooling multiple samples can address the limitations of low S/N, but a more fundamental problem remains for small inter-fluorophore distances: If the distance between two fluorescent spots is of the order of spot centre accuracy, the observed (Euclidean) distance systematically over-estimates the true distance, [5]. This over-estimation, or inflation, is a consequence of distances being convex functions (see Jensen’s inequality, [3, Thm 3.1.3]) and can be understood as a consequence of Euclidean distances having spherical level sets, Fig 1. The Euclidean distance is thus an inconsistent biased estimator. This bias also impacts polygon shape, for instance measured triangles become more equilateral (internal angles are biased towards 60°).

**Fig 1.**
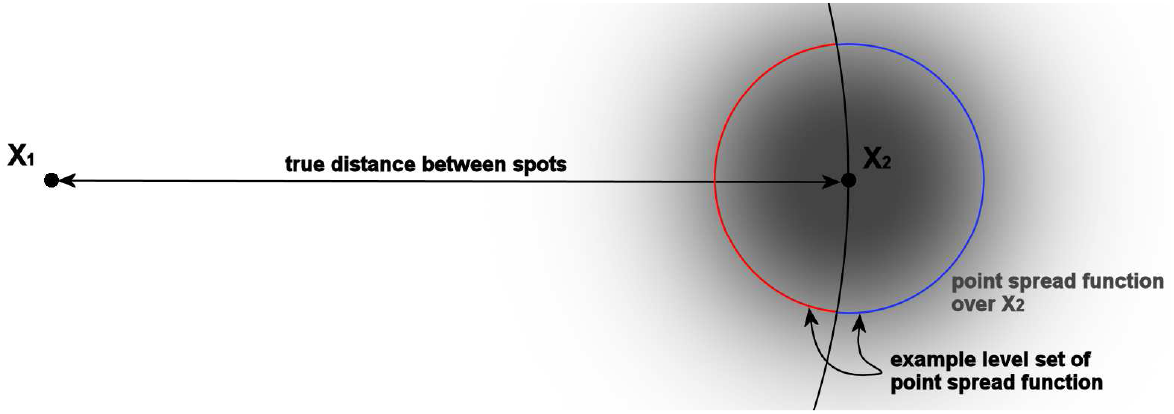
Schematic of the inflation of distances with diffraction limited spot measurements. The blue and red arcs together form a level set of the point spread function of spot 2 at true position *X*_2_ (grey). Thus the observed position of spot 2 is equally likely on any position on these arcs. The blue arc being longer than the red arc implies that spot 2 is more likely to be observed further away from spot 1 than their true distance. On average the observed distance is thus larger than the true distance.

The over-estimation of pair-wise distances is well understood, with computational correction methods being available, [5], [6], [13]. In particular, in the case of isotropic Gaussian measurement errors the likelihood is analytically tractable in 1-3 dimensions, [5], [6], and has been used across a variety of applications, [20], [10]. In 3D, however, typically PSFs are anisotropic, with the resolution along the optical axis usually reduced relative to that in the focal plane. For anisotropic measurement errors no analytical solution is known; however a Bayesian approach was developed for pair-wise distance correction [24]. Here we extend this methodology to multiple fluorophores/arbitrary polygons. Specifically we develop a Bayesian sampling algorithm (using a Markov Chain Monte Carlo (MCMC) framework) to infer a fixed polygon (referred to as the template) from observed samples, the polygon nodes being marked with distinct fluorophores and each sample rotated and translated relative to the template, Fig 2. This polygon based method has several benefits over pair-wise methods. Firstly, the geometric constraints are automatically satisfied in our model, e.g. for three fluorophores the triangle inequality holds. This contrasts to inferring the three lengths independently in a pair-wise fashion, where the inferred lengths do not necessarily make a triangle. Secondly, geometric correlations between the lengths can improve individual length confidence (reducing posterior variances). Thirdly, individual localisation errors of each fluorophore are inferred; this contrasts to pair-wise analyses where only the error of the displacement can be inferred.

**Fig 2.**
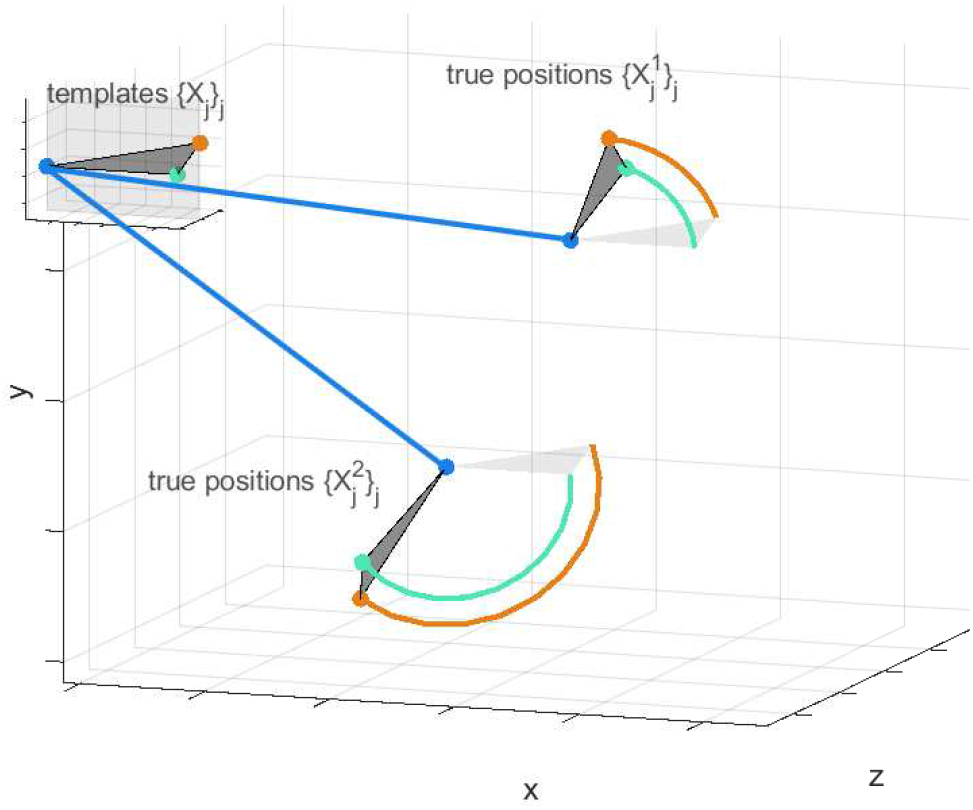
Template and sample perspectives. Inferred true fluo-rophore positions 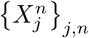 and template positions {*X*_*j*_}_*j*_ for *J* = 3 fluorophores and *N* = 2 measurements are shown. Each sample polygon is generated from the template by a translation and rotation (straight and circular lines). The observed positions 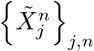 are omitted for simplicity.

We also extend the algorithm to enable analysis of heterogeneous datasets, i.e. samples from a population that comprise multiple polygon states. This allows us to study conformation changes of macro-molecular complexes *in situ*, our method being distinct from proximity sensors such as Förster Resonance Energy Transfer (FRET, [15]). We apply a two-state version of our algorithm to human kinetochores, macro-molecular complexes that play a vital role during cell division. Kinetochore attachment to microtubules is critical for the correct positioning of chromosomes in mitosis, microtubule attachment in fact leading to a conformational change within the kinetochore complex, [24]. By using our two-state model on triple-labelled kinetochores (re-analysing datasets from [24] thereby improving the resolution of those results), we demonstrate for the first time that there is conformational heterogeneity of the kinetochore complex during metaphase. We infer the sub-population size during metaphase of the unattached conformation.

This paper is organised as follows. Section Materials and methods presents the model within a Bayesian framework. Section Sampling based inference of model parameters: a Markov Chain Monte Carlo algorithm gives an outline of the inference algorithm. Full details of the algorithms can be found in the Supplementary Data Markov Chain Monte Carlo samplers for parameter inference. We demonstrate the accuracy of our algorithm on simulated/synthetic data in section Algorithm performance on simulated data, accurately inferring polygon side lengths, internal angles, the fluorophore measurement errors and - in the mixture analysis - the proportions, with which two states contribute to the mixture. For the single-state method we demonstrate the algorithm’s advantages over existing pair-wise analyses for triangle (three fluorophore) datasets. In section Analysis of the human kinetochore: a structured macro-molecular complex we use our algorithm to analyse the architecture of the human kinetochore from experimental 3D fluorescence imaging data, [24]. In subsection Experimental data for two-state mixture model we present results for the analysis of heterogeneous experimental samples assumed to comprise two states. In Conclusions we outline improvements and the limitations of our current algorithm.

## Materials and methods

To highlight the model’s generality, the model is presented for an arbitrary number *J* of distinct fluorophores. We later restrict our analysis to examples with three markers (*J* = 3, referred to as *triangle correction*). We initially assume that there is a single polygonal state, and then extend this model to mixtures of states in subsection Mixture of multiple states.

We consider fixed three-dimensional fluorescent images labelled with a number *J* of distinct markers that mark different sites within a macro-molecular structure. Here a site refers to part of the structure where the respective fluorescent labels cluster. In practice, each marker should be sufficiently tightly localised for the Gaussian spot approximation to be valid, with a width similar to a diffraction limited object. Fluorescent labels can be fluorescently labelled antibodies, labelled DNA (FISH), or a genetically encoded fluorophore such as GFP. The objective is to infer the underlying polygon (size, edge lengths, angles between edges) from a sample of observed marker positions, where the *J* markers are located at the nodes of the polygon. The true marker positions are unknown because of measurement noise.

### Model assumptions

We assume a sample of *N* measurements of the *J* sided polygon. The location and orientation (here jointly referred to as the *perspective*) of this polygon are specific to each measurement. Thus, there is a translation and 3D rotation associated with each measured polygon, see Fig 2. We assume space is uniform, so the (true) polygons are uniformly distributed throughout space and undergo an isotropic rotation.

We assume that the measurement errors are anisotropic Gaussians with covariance matrix diag 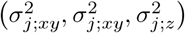 in the given coordinate system (*z* along the optical axis) for marker type *j* ∈ {1, … , *J*}; previous pair-wise methods also assume Gaussian errors, [5] (isotropic) and [13] (anisotropic). Note, that only the shape of the measurement error is assumed; *σ*_*j*;*xy*_, *σ*_*j*;*z*_ are to be inferred. Errors are fluorophore specific because quantum efficacy differs between fluorophores (photon count emission), and may also depend on the imaging conditions and the number of labelled fluorophores at a site. Typical data consists of spot centres determined by fitting 3D Gaussian profiles to spots in the image (either individually or using a mixture of Gaussians model).

### Polygon parametrisation and inference

The underlying polygon is defined by a standardised template, with nodes at positions *X*_*j*_ ∈ ℝ^3^. In the following we refer to this as the template, and the nodes as the template positions. We write {*X*_*j*_}_*j*_ to denote the set of all template positions ({*X*_*j*_}_*j*∈{1,…,*J*}_). We use a similar notation for other parameters. We choose the template such that *X*_1_ is at the origin, *X*_2_ is on the non-negative *x*-axis and *X*_3_ is in the *x*-*y*-plane with non-negative *y*. Assume there are *N* observations of this polygon, each associated with a perspective - the perspective for a particular measurement *n* ∈ {1, … . *N*} relative to the template is defined by six parameters: the first three define a translation *T*^*n*^ ∈ ℝ^3^ of the true position of marker *j* = 1 relative to *X*_1_; the next three are the Euler angles of the three-dimensional rotation *R*^*n*^ ∈ *SO* (3) of the true marker position relative to the template, with centre of rotation at marker *j* = 1, Fig. 2. Thus, the true positions of the markers of each measurement are given by:

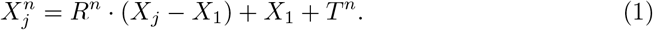

The observed marker position 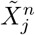 incorporates (Gaussian) measurement error 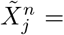 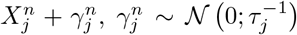 independently for all *j* and *n*, with (3×3) precision matrix 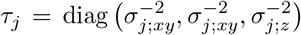. The likelihood for the model parameters {*θ*_*j*_}_*j*_ ≔ {{*X*_*j*_}_*j*_, {*τ*_*j*_}_*j*_}, *j* ∈ {1, …, *J*}, and perspectives {*ϑ*^*n*^}_*n*_ ≔ {{*T*^*n*^}_*n*_, {*R*^*n*^}_*n*_}, *n* over the set of samples {1, …, *N*}, reads:

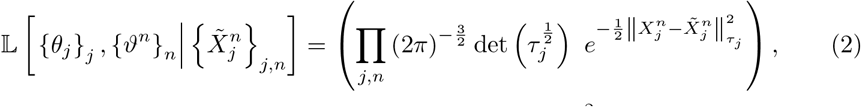

where index ranges are suppressed for simplicity, and 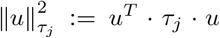 denotes the squared Euclidean norm of vector *u* ∈ ℝ^3^, weighted with *τ*_*j*_ (here *T* denotes the transpose). The predicted true positions 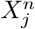 for marker *j*, measurement *n*, is given in Eq (1). The likelihood thus has a dependence on the hidden perspective variables *T*^*n*^, *R*^*n*^.

The posterior *dπ* is given, up to proportionality, by multiplying the likelihood from Eq (2) with the prior, denoted *dπ*_0_ [{*θ*_*j*_}_*j*_, {*ϑ*^*n*^}_*n*_]. We use uninformative priors, namely translations are homogeneously distributed in space, rotations are isotropic, measurement errors are flat on the positive half-space and the prior on the template positions is (approximately) flat on the marginal of each polygon length |*X*_*j*_ − *X*_*i*_|. For details, see section Uninformative priors.

### Mixture of multiple states

The above model assumes population homogeneity, *i.e.* there is only a single underlying polygon state, positions {*X*_*j*_}_*j*_, from which all observed polygons, 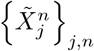, arise (up to measurement noise and perspective). This is a strong assumption, and may be violated in applications. Specifically there may be a mixture of underlying polygon states with distinct polygon templates and measurement errors for each state. Here we extend the model to allow for heterogeneity of the (polygon) state; i.e. measurements originate from one of *Z* polygon templates 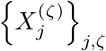 for *ζ* ∈ {1, … . *Z*}. In addition, the corresponding measurement errors may vary between the states, *i.e.* we have

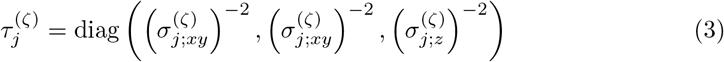

extending the notation above. In this case we not only want to infer the template positions 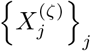 and measurement errors 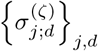 for each state *ζ*, but also the proportion *p*^(*ζ*)^ ∈ [0, 1] of each state in the mixture. In the application we confine ourselves to *Z* = 2.

Analogous to Eq (1) the predicted true positions are in this case:

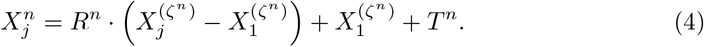

Note that the marker positions of the *n*^*th*^ measurement derive from the template *ζ*^*n*^. The state-affiliations {*ζ*^*n*^}_*n*_ are hidden variables, which need to be inferred. We use a generalised Bernoulli-distributed prior (also known as the multinoulli or categorical distribution) on *Z* categories for the states, *i.e.* 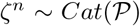 independently for each *n*, where 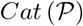 denotes the categorical distribution with parameters 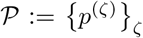, the probabilities of the respective states, *ζ* ∈ {1, … , *Z*} (hence *p*(*ζ*) ∈ [0, 1] and (Σ_*ζ*∈{1,…,*Z*}_*p*^(*ζ*)^) = 1).

We use a Dirichlet distributed (hyper-) prior:

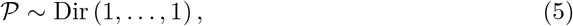

i.e. there is no prior preference for any state in the mixture, (see Eq (S3.10) for the distribution).

Thus, the likelihood function of the extended model reads:

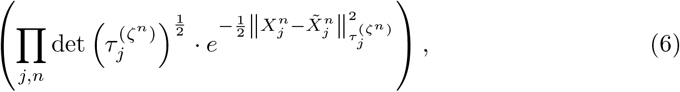

where 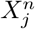 is the state-dependent Eq (4), while the prior of the extended model is:

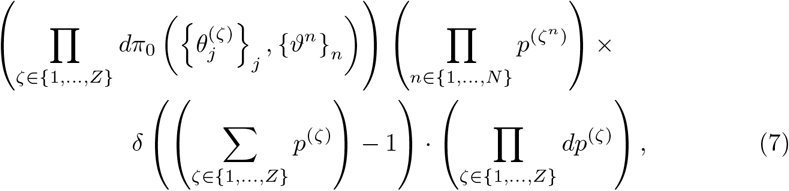

where *p*^(*ζ*)^ are valued in [0, 1], and 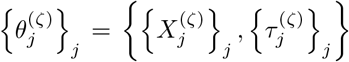. Note, that this reduces to the basic single-state version described before when *Z* = 1.

In practice, we found for mixture models that uninformative priors, Eq (7), can result in poor convergence. There were two issues. Firstly, if a state is “lost” during a step in the Markov chain, (i.e. state *ζ*′ does not occur, ∀_*n*_ : *ζ*^*n*^ ≠ *ζ*′), then its reappearance can require a substantial number of steps. For that reason, we impose that each state has to occur at least three times (i.e. 3 ≤ |{*n* ∈ {1, … , *N*}|*ζ*^*n*^ = *ζ*′}| for all *ζ*′ ∈ {1, … , *Z*). In all examples considered here, the posterior was far away from this boundary. The second issue is when a polygon state is rare, then both the posterior shape/size of the rare state polygon, and its proportion *p*^(2)^ will have low confidence (assuming the rare state is *ζ* = 2). To improve posterior proportion confidence we used a joint inference methodology, utilising two datasets where the second dataset was assumed to comprise homogeneous (pure) samples of the *ζ* = 2 state. By utilising samples to define the pure state, we capture all the correlations of that state, which would be hard to define through priors. This purity condition could be relaxed, we only require that a state occurs at a significant fraction in at least one dataset. We call the respective dataset the *state-informing dataset*. The above constraint on each state occurring at least three times is imposed across the datasets and thus trivially satisfied. The likelihood for joint inference is the product of the likelihoods for each dataset, either the multi-state likelihood for mixed datasets or the single-state likelihood for the pure datasets.

On these joint inference models, we sometimes utilised box priors on one or more of the states to improve convergence. Specifically, the triangle side lengths are independently constrained to a box and the precisions are Gamma distributed. For a box prior on state *ζ* = 1, the following factors are included in the prior,

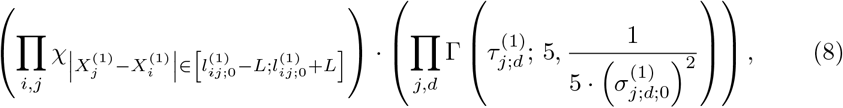

where the parameters 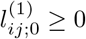 and 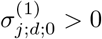 are specified by prior knowledge (analysis of previous data or structural data), and the box size 2*L* is taken as 24nm. The second and third parameter of the Γ distribution are the shape and scale parameters. The values for *L* and the shape parameter are a particular choice for our examples.

### Sampling based inference of model parameters: a Markov Chain Monte Carlo algorithm

To infer the model parameters - the template and measurement errors {*θ*_*j*_}_*j*_ and the sample specific perspectives {*ϑ*^*n*^}_*n*_ (and for the multi-state model, the state affiliations {*ζ*^*n*^}_*n*_ and proportions {*p*^*ζ*^}_*ζ*_ as well) - we use a Bayesian computational method. Specifically, the posterior distribution of the parameters given the data is, up to proportionality (for the single state model)

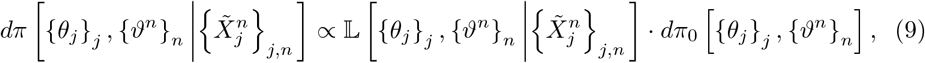

with the likelihood from Eq (2) and prior *π*_0_ from Eq (S1.1). There are a range of algorithms that can be used to sample from this posterior. We use a Markov Chain Monte Carlo (MCMC) methodology, [14, Ch 1], [18, Ch 1], whereby a Markov chain is constructed that has a stationary distribution equal to the posterior distribution. Once converged, the chain can be used to sample from the posterior.

In the single-state model we sequentially update each of the parameters separately: the perspectives, {*R*^*n*^}_*n*_, {*T*^*n*^}_*n*_, for each polygon sample *n*, the template positions, {*X*_*j*_}_*j*_, and the precisions, {*τ*_*j*_}_*j*_, for each marker *j*, respectively. For {*X*_*j*_}_*j*_, {*R*^*n*^}_*n*_ and {*T*^*n*^}_*n*_, random walk samplers are used, while for {*T*^*n*^}_*n*_ and {*τ*_*j*_}_*j*_ we use Gibbs samplers.

In the multi-state model we sequentially update the perspectives just as in the single-state model, while each template position 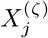 and precision 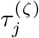 is updated sequentially for each marker *j* and state *ζ*. The state proportions {*p*^(*ζ*)^}_*ζ*_ are updated jointly to always satisfy their normalisation condition. For each of the 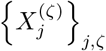, {*R*^*n*^}_*n*_, {*T*^*n*^}_*n*_ and {*ζ*^*n*^}_*n*_ random walk samplers are used, while for {*T*^*n*^}_*n*_, 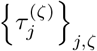 and {*p*^(*ζ*)^}_*ζ*_ we use Gibbs samplers. See subsection Markov Chain Monte Carlo samplers for parameter inference for algorithm details.

## Results

### Algorithm performance on simulated data

We test our polygon inference algorithm on simulated data for both the single-state and the multi-state models, confirming that our algorithms reproduce the true original parameters of the simulations. For the single-state model there are existing methods for length correction: the pair-wise Bayesian Euclidean Distance Correction Algorithm (BEDCA, [13]), and the analytic length correction based on [5]. The latter is only applicable to isotropic measurement errors and comes in two variations, a maximum likelihood estimate developed in [5] (“MLE” below) and a full posterior probability version (“means” below; see Implementation of pair-wise correction methods). We compare our triangle algorithm to these methods (where applicable). Note we use a flat prior on the measurement errors in all methods for fair comparison. We confine ourselves to *J* = 3 markers, referring to our method as the triangle correction (method). For the multi-state model we use *Z* = 2 states.

#### Testing the single-state model on simulated data

We illustrate our algorithm on simulated data with (true) triangle lengths and measurement errors similar to those observed in biological complexes, [24]. We simulate data with *J* = 3 markers and *N* = 400 independent measurements, using the core model described in subsection Polygon parametrisation and inference, see supplementary section Simulated data for details. True parameters and the corresponding inferred values are shown in Table 1 for six examples; see Fig 3 for the posterior distributions of the inferred lengths and internal angles. Our correction algorithm typically gives inferred values that are in good agreement with the true values. These examples clearly confirm that the Euclidean distance estimator 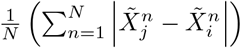 is subject to significant length inflation.

**Table 1.** Single-state simulated examples (N = 400). Rows are the original (true) value, and the posterior means (or MLE) standard deviations in subsequent rows for stated method. ‘Direct measurement’ refers to the (uncorrected) Euclidean distance 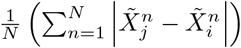, which is known to overestimate distances [6]. “Churchman means” refers to the posterior mean using a flat prior on the length and measurement error, *dl*_*ij*_*dσ*_*ij*_. We quote the pair-wise variances to compare to the pair-wise algorithm (for triangle we have individual fluorophore measurement errors to give pair-wise variance using 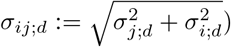. For inferred posterior distributions of the lengths see Fig 3.

| (lengths and errors in nm) | $\vert X_2 - X_1\vert $ | $\vert X_3 - X_1\vert $ | $\vert X_3 - X_2\vert $ | $\sigma_{12;xy}$ | $\sigma_{12;z}$ | $\sigma_{13;xy}$ | $\sigma_{13;z}$ | $\sigma_{23;xy}$ | $\sigma_{23;z}$ |
| --- | --- | --- | --- | --- | --- | --- | --- | --- | --- |
| Example 1 with isotropic measurement error: |  |  |  |  |  |  |  |  |  |
| true value | 50 | 60 | 15 | 21.2 | 21.2 | 29.2 | 29.2 | 29.2 | 29.2 |
| direct measurement | $59.1 \pm 1.0$ | $73.2 \pm 1.3$ | $49.2 \pm 1.0$ | / | / | / | / | / | / |
| Churchman MLE, [5] | $50.5 \pm 1.5$ | $58.9 \pm 2.5$ | $33.2 \pm 3.5$ | $20.9 \pm 0.9$ | | $29.3 \pm 1.5$ | | $23.7 \pm 1.6$ | |
| Churchman means, [5] | $50.2 \pm 1.5$ | $58.2 \pm 2.7$ | $27.0 \pm 9.3$ | $21.1 \pm 1.0$ | | $29.7 \pm 1.6$ | | $25.6 \pm 2.6$ | |
| BEDCA, [13] | $50.2 \pm 1.6$ | $58.0 \pm 2.8$ | $27.1 \pm 9.2$ | $21.2 \pm 1.2$ | $21.0 \pm 1.5$ | $30.2 \pm 1.9$ | $29.2 \pm 2.1$ | $25.4 \pm 2.6$ | $26.0 \pm 2.9$ |
| triangle, uninformative prior | $51.8 \pm 2.1$ | $58.2 \pm 3.6$ | $12.3 \pm 6.7$ | $20.3 \pm 1.5$ | $23.0 \pm 2.2$ | $31.3 \pm 2.6$ | $29.2 \pm 2.7$ | $30.5 \pm 1.6$ | $31.9 \pm 1.9$ |
| Example 2 with anisotropic measurement error: |  |  |  |  |  |  |  |  |  |
| true value | 50 | 60 | 15 | 14.1 | 28.3 | 18.0 | 36.1 | 18.0 | 36.1 |
| direct measurement | $58.2 \pm 0.9$ | $70.0 \pm 1.2$ | $43.3 \pm 1.0$ | / | / | / | / | / | / |
| BEDCA, [13] | $50.6 \pm 1.2$ | $60.5 \pm 1.4$ | $16.1 \pm 7.0$ | $14.0 \pm 0.9$ | $28.7 \pm 1.6$ | $17.7 \pm 1.2$ | $36.1 \pm 2.0$ | $17.2 \pm 1.9$ | $36.7 \pm 1.7$ |
| triangle, uninformative prior | $50.9 \pm 1.1$ | $60.4 \pm 1.5$ | $14.7 \pm 3.0$ | $13.6 \pm 0.8$ | $29.3 \pm 1.6$ | $18.3 \pm 1.2$ | $34.7 \pm 1.9$ | $18.1 \pm 0.9$ | $37.2 \pm 1.4$ |
| Example 3 with anisotropic measurement error: |  |  |  |  |  |  |  |  |  |
| true value | 45 | 60 | 15 | 14.1 | 28.3 | 18.0 | 36.1 | 18.0 | 36.1 |
| BEDCA, [13] | $45.9 \pm 1.2$ | $59.9 \pm 1.4$ | $12.2 \pm 6.4$ | $14.4 \pm 0.9$ | $27.8 \pm 1.6$ | $17.5 \pm 1.0$ | $34.3 \pm 2.0$ | $17.3 \pm 1.5$ | $35.5 \pm 1.5$ |
| triangle, uninformative prior | $46.2 \pm 1.1$ | $59.8 \pm 1.4$ | $15.4 \pm 1.8$ | $14.2 \pm 0.8$ | $27.7 \pm 1.5$ | $17.4 \pm 1.0$ | $34.9 \pm 1.9$ | $16.9 \pm 0.6$ | $35.3 \pm 1.3$ |
| Example 4 with anisotropic measurement error: |  |  |  |  |  |  |  |  |  |
| true value | 60 | 60 | 15 | 14.1 | 28.3 | 18.0 | 36.1 | 18.0 | 36.1 |
| BEDCA, [13] | $58.2 \pm 1.0$ | $57.9 \pm 1.4$ | $13.4 \pm 6.7$ | $13.0 \pm 0.8$ | $27.2 \pm 1.6$ | $16.5 \pm 1.1$ | $38.6 \pm 2.1$ | $17.1 \pm 1.7$ | $38.6 \pm 1.6$ |
| triangle, uninformative prior | $58.4 \pm 1.0$ | $57.4 \pm 1.4$ | $15.1 \pm 3.9$ | $12.8 \pm 0.8$ | $27.5 \pm 1.5$ | $17.2 \pm 1.1$ | $38.8 \pm 1.8$ | $16.9 \pm 1.1$ | $38.7 \pm 1.5$ |
| Example 5 with anisotropic measurement error: |  |  |  |  |  |  |  |  |  |
| true value | 15 | 15 | 15 | 14.1 | 28.3 | 18.0 | 36.1 | 18.0 | 36.1 |
| BEDCA, [13] | $16.4 \pm 5.3$ | $18.8 \pm 6.8$ | $9.08 \pm 5.42$ | $13.6 \pm 1.7$ | $29.0 \pm 1.4$ | $16.5 \pm 2.1$ | $35.7 \pm 1.7$ | $19.4 \pm 1.1$ | $37.3 \pm 1.4$ |
| triangle, uninformative prior | $17.5 \pm 4.4$ | $17.8 \pm 4.4$ | $7.77 \pm 5.15$ | $13.4 \pm 1.5$ | $28.8 \pm 1.4$ | $17.2 \pm 1.3$ | $36.1 \pm 1.5$ | $19.7 \pm 1.0$ | $37.5 \pm 1.4$ |
| Example 6 with anisotropic measurement error: |  |  |  |  |  |  |  |  |  |
| true value | 60 | 60 | 0 | 14.1 | 28.3 | 18.0 | 36.1 | 18.0 | 36.1 |
| BEDCA, [13] | $60.5 \pm 1.1$ | $60.9 \pm 1.3$ | $10.1 \pm 5.8$ | $14.0 \pm 0.9$ | $28.1 \pm 1.7$ | $17.0 \pm 1.0$ | $34.8 \pm 1.9$ | $16.9 \pm 1.3$ | $35.8 \pm 1.4$ |
| triangle, uninformative prior | $60.6 \pm 1.0$ | $60.9 \pm 1.1$ | $3.5 \pm 2.9$ | $13.9 \pm 0.8$ | $28.2 \pm 1.6$ | $17.6 \pm 0.9$ | $33.1 \pm 1.7$ | $18.1 \pm 0.5$ | $36.4 \pm 1.3$ |

**Fig 3.**
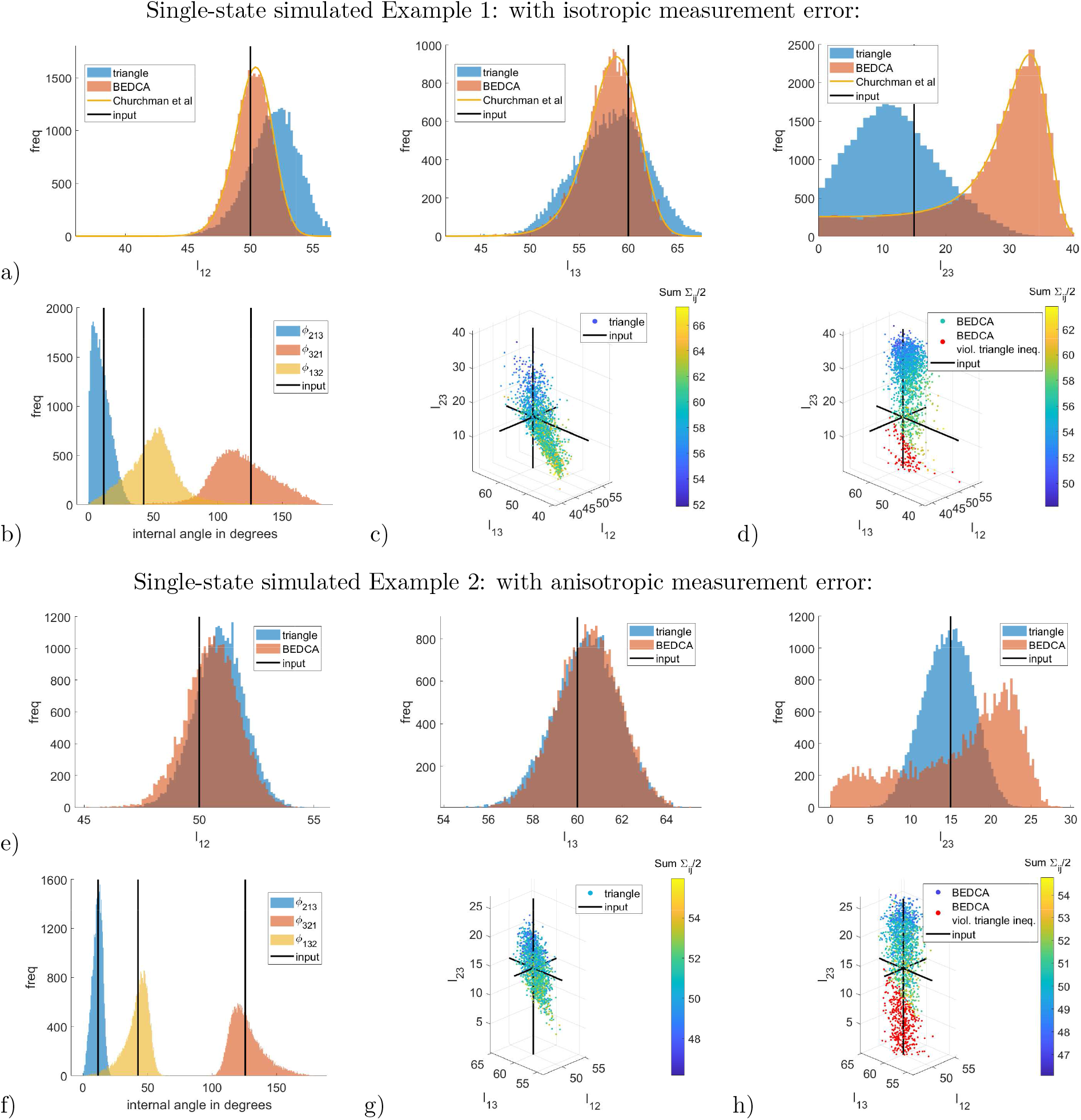
Posterior distributions for single-state simulated Examples 1,2. Marginal inferred posteriors for lengths and internal angles. For each example the top row (a,e) shows the marginal posteriors of the three triangle lengths as inferred by each of the correction methods. The bottom left images (b,f) show the inferred internal angles using the triangle correction presented here. The bottom centre images (c,g) show a scatter plot of the joint distribution as obtained with the triangle correction. The bottom right images (d,h) show the naive attempt to achieve a joint distribution of the three triangle lengths from the BEDCA method [13] by assuming independence. Samples that violate the triangle inequality are shown in red (about 7* of the samples in Example 1, 22* in Example 2), otherwise the colour represents the sum of all measurement errors, specifically 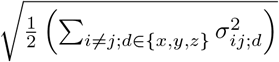 , to indicate the correlation between inferred lengths and errors. The plots show random sub-samples of the total sample size, to improve visibility. See Table 1 for the posterior means and standard deviations. See section Additional supplementary images for the corresponding Markov chain evolution of the parameters of the triangle inference.

All correction methods have similar performance for sufficiently large lengths (relative to the measurement error) and isotropic (Gaussian) measurement errors, Example 1, sides |*X*_2_ − *X*_1_| , |*X*_3_ − *X*_1_|. For small lengths, less or similar to the measurement error, all correction methods suffer from relatively large inference errors, length |*X*_3_ − *X*_2_| in our example with true value 15nm, (see Small lengths are increasingly difficult to infer for an explanation). Problems with the maximum likelihood estimate from [5] (“MLE” in Table 1) were reported previously, [20], with results becoming error-prone for lengths similar to, or smaller than the measurement error. For these short lengths we find this method tends to report misleading results with overconfident error-estimates (Example 1 in Table 1). For pair-wise inferences the likelihood with isotropic measurement errors is analytically tractable (Eq (6) in [5]); we can thus determine the posterior (assuming the same flat priors we use for the pair-wise method) and therefore calculate the posterior mean (“means” in Table 1). This estimate, and the pair-wise based Bayesian inference method [13] give near identical results (as expected as they share the same assumptions apart from the extra degree of freedom of [13] for anisotropic measurement errors). For the small length |*X*_3_ − *X*_2_|, the pair-wise based posterior has a long thick tail towards zero, see Fig 3a,c, indicating large uncertainty. The triangle correction has a posterior mean much closer to the true value and has substantially smaller variance, Table 1, Example 1. This is because information is essentially shared between the three lengths, allowing inference of the smaller length to be improved. There is an associated increase in the uncertainty of the other two lengths relative to the pair-wise correction, Table 1, Example 1.

Examples 2-6 in Table 1 are for anisotropic measurement errors. There are only two methods available for anisotropic errors: our triangle correction and the pair-wise Bayesian method [13]. These two methods give consistent results, but the triangle method typically has lower posterior variance, particularly for the short lengths. For a true length of zero, Example 6, the triangle method is substantially better (provided the other two lengths are large). The triangle correction improves on pair-wise based inference in the following ways:

- It can infer small lengths with higher confidence, Examples 1–6. A triplet of fluorophores can thus be utilised to improve inference of a small distance by placing a third fluorophore distant from the two fluorophores of interest (50-200nm). This is because correlations between the lengths confer information on the smallest length. This is demonstrated for the small length |*X*_3_ − *X*_2_| in Examples 2–6, where the examples with the more distant auxiliary marker *j* = 1 and the more co-linear triangle geometries exhibit the largest benefits.
- It infers the triangle (more generally the polygon) and therefore reconstructs the entire geometry of the markers, including the internal angles. Reconstructing polygons from pair-wise estimates, assuming independence of the inferred lengths, can lead to violations of the triangle inequality (see scatterplots in Fig 3), and more generally polygon based constraints.
- Using three or more fluorophores allows measurement errors to be inferred individually for each marker. For pair-wise methods these are not accessible, as only the error of the displacement *X*_*j*_ − *X*_*i*_ can be estimated. This follows since the system of equations 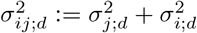 for *j* ≠ *i* ∈ {1, … , *J*} can be solved uniquely for each 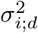 for *J* = 3, but not for *J* = 2 (there are *J* unknowns for 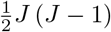 constraints). See Table S14.1 for marginal posteriors parameters of Examples 1, 2.

#### Testing the two-state model on simulated data

To test the multi-state mixture algorithm, we determined inference accuracy of the states and composition of a mixture of *Z* = 2 triangle states on simulated triangle data (*J* = 3). We demonstrate joint inference from a mixed dataset and a pure dataset. The first has a mixture of the two states, *N*_mix_ = 600 measurements (*Z* = 2 states, *J* = 3 markers), with proportions *p*^(1)^ and *p*^(2)^ = 1 − *p*^(1)^ for the states *ζ* = 1, *ζ* = 2, where state *ζ* = 2 is the minor population (*p*^(2)^ ≤ *p*^(1)^). This data is simulated from the multi-state model of subsection Mixture of multiple states (see supplementary section Simulated data for simulation details). The second dataset is a pure population comprising only state *ζ* = 2. We simulate *N*_inform_ = *p*^(1)^ · *N*_mix_ from the single state model for the pure dataset. Thus, there are the same number of samples in the pure dataset as samples of the dominant *ζ* = 1 state in the mixed dataset.

We impose an additional prior on the *ζ* = 1 state to limit exploration of the parameter space during burnin; this reduces the convergence time. Specifically, we use the prior from Eq (8), i.e. a box prior on the triangle side lengths and gamma-distributed priors on the measurement errors. The parameters 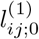 and 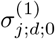 are chosen so the means of the priors are equal to the true values of the simulated data. We confirmed that the boxes are sufficiently large so that the prior has negligible impact on the posterior. Specifically, the bulk of the marginal length posteriors are well within the boundaries of their box (in fact, in all our examples, the weight of a Gaussian fit to the joint posterior of the three lengths that was outside the box never exceeded 5*). However, the priors on the measurement errors are not weak and can increase confidence on less informative datasets. The effect of this prior on the posterior (of the experimental examples studied in subsection Experimental data for two-state mixture model) is explored in supplement Effect of priors in two-state model.

Simulated examples and the corresponding inferred parameters (triangle lengths and state proportions) are summarised in Table 2. The (marginal) posteriors for the state proportion *p*^(2)^ are shown in Fig 4, demonstrating clear localisation of the posterior around the true value with posterior standard deviations of the order of 6* (for these sample sizes and parameter values). Thus, the state proportion posteriors are only well separated from zero for true proportions above 10*; for 20* and above there is clear evidence for the presence of the minor population. All lengths are inferred correctly, except the true values of two lengths in Example 2 lie in the posterior tail. Since we are inferring 42 parameters in Table 2, it is expected that some deviations will occur. Reruns of Example 2 on new datasets gave posteriors consistent with the true values. Inferred measurement errors (omitted for brevity) were all consistent with the true values (for both 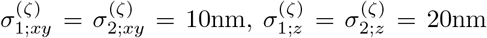 and 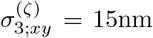, 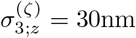).

**Table 2.** Two-state simulated examples (anisotropic measurement error). Rows are the true values and posterior means and standard deviations for respective examples, columns the two triangle states (lengths) and the state proportion *p*^(2)^ in the mixture. Note, Example 2 has posterior means significantly different from the true value by more than two standard deviations, denoted by * (*p* = 0.8* and *p* = 0.2* for the inferred versus the true values of 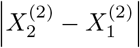 and 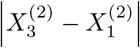, respectively).

| (lengths and errors in nm) | $\vert X_2^{(1)} - X_1^{(1)}\vert $ | $\vert X_3^{(1)} - X_1^{(1)}\vert $ | $\vert X_3^{(1)} - X_2^{(1)}\vert $ | $\vert X_2^{(2)} - X_1^{(2)}\vert $ | $\vert X_3^{(2)} - X_1^{(2)}\vert $ | $\vert X_3^{(2)} - X_2^{(2)}\vert $ | $p^{(2)}$ |
| --- | --- | --- | --- | --- | --- | --- | --- |
| Example 1: |  |  |  |  |  |  |  |
| true value | 45 | 85 | 45 | 50 | 60 | 15 | 0% |
| triangle, Eq (8) prior | $45.7 \pm 0.9$ | $85.3 \pm 1.1$ | $45.1 \pm 1.3$ | $50.7 \pm 0.9$ | $60.6 \pm 1.0$ | $17.1 \pm 2.1$ | $(1.99 \pm 1.74) \%$ |
| Example 2: |  |  |  |  |  |  |  |
| true value | 45 | 85 | 45 | 50 | 60 | 15 | 10% |
| triangle, Eq (8) prior | $46.1 \pm 1.0$ | $85.6 \pm 1.7$ | $44.8 \pm 2.1$ | $52.0 \pm 0.8^*$ | $63.5 \pm 1.1^*$ | $18.9 \pm 2.1$ | $(9.95 \pm 5.45) \%$ |
| Example 3: |  |  |  |  |  |  |  |
| true value | 45 | 85 | 45 | 50 | 60 | 15 | 20% |
| triangle, Eq (8) prior | $46.0 \pm 1.1$ | $83.3 \pm 2.3$ | $41.7 \pm 3.3$ | $51.0 \pm 0.9$ | $61.3 \pm 1.2$ | $18.1 \pm 2.1$ | $(18.4 \pm 7.1) \%$ |
| Example 4: |  |  |  |  |  |  |  |
| true value | 45 | 85 | 45 | 50 | 60 | 15 | 30% |
| triangle, Eq (8) prior | $43.7 \pm 1.4$ | $88.0 \pm 2.6$ | $48.0 \pm 3.3$ | $50.0 \pm 1.0$ | $59.8 \pm 1.3$ | $14.7 \pm 2.6$ | $(30.8 \pm 6.0) \%$ |
| Example 5: |  |  |  |  |  |  |  |
| true value | 45 | 85 | 45 | 50 | 60 | 15 | 40% |
| triangle, Eq (8) prior | $44.4 \pm 1.4$ | $89.1 \pm 2.9$ | $49.5 \pm 3.4$ | $49.8 \pm 1.1$ | $59.6 \pm 1.3$ | $14.1 \pm 2.7$ | $(40.3 \pm 6.0) \%$ |
| Example 6: |  |  |  |  |  |  |  |
| true value | 45 | 85 | 45 | 50 | 60 | 15 | 50% |
| triangle, Eq (8) prior | $45.0 \pm 1.6$ | $88.5 \pm 3.9$ | $48.0 \pm 4.8$ | $49.7 \pm 1.1$ | $59.8 \pm 1.4$ | $14.3 \pm 2.7$ | $(49.5 \pm 7.5) \%$ |

**Fig 4.**
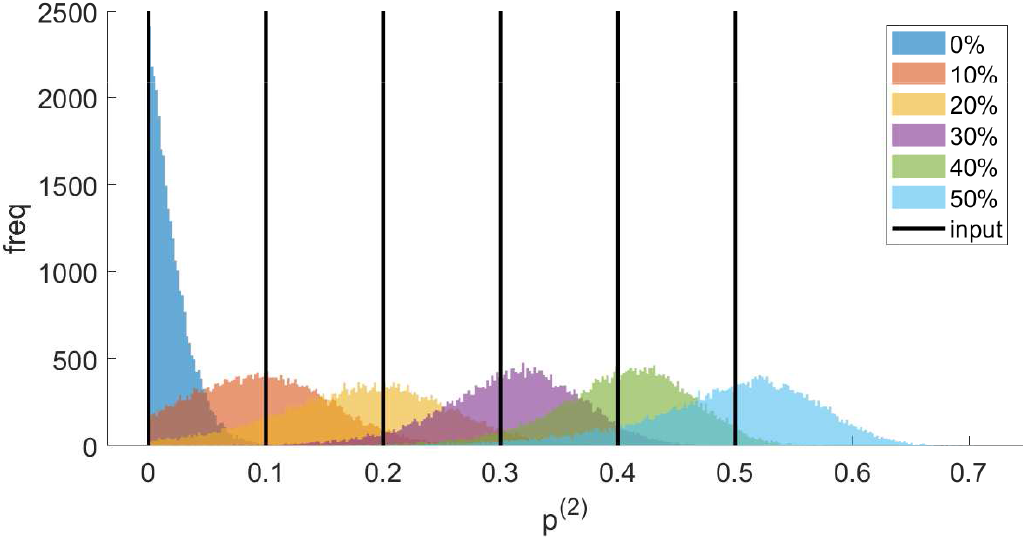
Two-state simulated examples. Posterior state proportions for the simulated two-state model with varying state proportions *p*^(2)^. Original (true) values are marked with black vertical lines, posteriors of each simulated dataset indicated in legend.

### Analysis of the human kinetochore: a structured macro-molecular complex

#### The experimental system: human cell division

To test performance of our algorithms on experimental data, we used three-fluorophore 3D images of human kinetochores from hTERT-immortalised retinal pigment epithelial (RPE) cells, imaged during metaphase of cell division with a confocal spinning-disk microscope, [24]. Kinetochores are a macro-molecular complex that orchestrate chromosome movements during cell division, playing a vital role in congression and segregation dynamics [22]. Kinetochores interface between the spindle and chromosomes, binding to both DNA (through the histone CenpA) and microtubules, and thus connecting chromosomes to the spindle machinery.

It is multi-functional, being a force generator and a sensor for erroneously connected microtubules [25]. During S-phase of cell division, chromosomes are duplicated– these duplicates are held together by condensins near the kinetochores, giving the familiar ‘X’ shape of mitotic chromosomes. These duplicated chromosomes are called sister chromatids, or simply sisters. At the start of mitosis (M-phase), after nuclear envelope breakdown, kinetochores are not bound to microtubules. As mitosis progresses, kinetochores attach to microtubules emanating from the spindle poles, and ideally sister kinetochores attach to opposite poles to form a bi-orientated state, Fig 5. Bi-orientated chromosomes congress to the cell mid-plane, in essence a holding configuration whilst any remaining chromosomes are captured and bi-orientated, Fig 5. Microtubule attachment is sensed by the kinetochore, feeding into the spindle assembly checkpoint (SAC), [25], that delays anaphase (segregation of chromosomes to daughter cells) until erroneous attachments are no longer detected. Once all kinetochores are bi-orientated the spindle check-point is satisfied and the cell enters anaphase, sister chromatids separating into distinct daughter cells.

**Fig 5.**
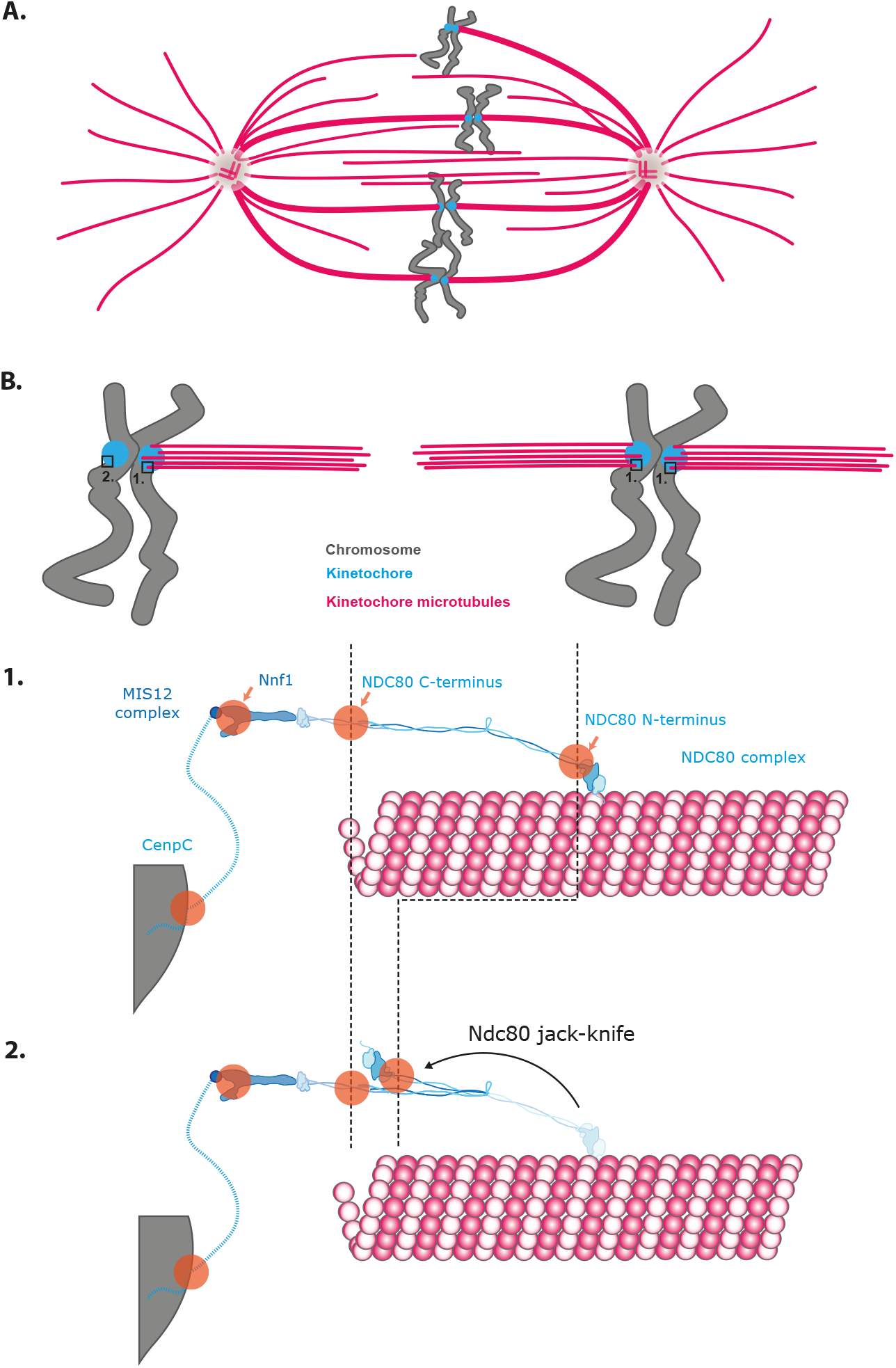
Spindle and kinetochore organisation. **A.** Schematic of the spindle with most chromatids congressed at the cell mid-plane and a late congressing chromatid. Chromatids are bi-orientated with microtubule attachments to both spindle poles, except the top chromatid pair which has only one kinetochore attached to the right spindle pole (termed monotelic). **B**. Detail of monotelic and bi-orientated sister pairs. The kinetochores are in states 1. (attached) or 2. (unattached), shown below. State 1. - attached to microtubules. Molecular detail of the Ndc80 complex showing its attachment to the side of a microtubule at its N-terminal binding site. The Ndc80 is attached to the Nnf1 and Mis12 complexes at its C-terminus, which binds CenpC through an unstructured linkage. Shown are the positions of the four fluorophores examined in this paper. State 2. - unattached Ndc80. The Ndc80 complex is hinged, and in the unattached state the Ndc80 complex jackknifes bringing the N and C termini closer together. A kinetochore has about 200–250 Ndc80 complexes and is bound to around 20 microtubules in mature attachment.

A key constituent of kinetochores is the Ndc80 complex that binds to microtubules, Fig 5B. The Ndc80 complex has a hinge about 16nm from the N terminus, which allows the complex to be either fully straightened or folded back (jackknifed). This conformational change is observed upon microtubule binding, [24]; specifically, the microtubule binding site (N terminus, denoted Ndc80N) moved ≈ 25nm within the kinetochore between microtubule binding conditions (attached) and depolymerised microtubule conditions (unattached), Fig 5B, states 1. and 2. The Ndc80 complex thus goes from a straightened configuration when attached, to jackknifed, or folded, when unattached. Relative to a 3rd fluorophore, we thus have two distinct triangular states. These triangles were reconstructed from a pair-wise analysis in [24]. By analysing three-fluorophore data with our triangle correction algorithm, the triangle states associated with attached and unattached can be determined.

The kinetochore comprises multiple copies of constituent proteins; there are an estimated 200–250 Ndc80 molecules ( [28], HeLa cells) and it attaches a bundle of around 20 microtubules [2]. As discussed in [24], our results refer to this ensemble, which is a diffraction-limited spot in each channel. However, the kinetochore is highly structured. This high ordering within the kinetochore means that the changes in the triangular state reflect conformational changes of the Ndc80 molecule itself.

As cell division progresses and each sister chromatid pair achieves bi-orientation, the cell’s 46 kinetochores change from unattached (in prometaphase) to all attached at the end of metaphase, immediately before anaphase. Hence, there is expected to be a minor proportion of kinetochores in metaphase that are still in the unattached state, *i.e.* the population is heterogeneous. However, there is no direct evidence of this conformation change occurring during cell division, or confirmation that the kinetochore population is conformationally heterogeneous. The presence of unattached kinetochores can be inferred by observing if check-point proteins (such as Mad, Bub) are recruited, [7]. However, the check-point is a downstream integration of attachment and inter-sister tension signals, [27], so is only an indirect indicator of attachment state. Here, we use our mixture model to determine if we can detect that minor population in the unattached conformational state and to estimate its proportion.

#### Experimental data for single-state model

We consider five examples of triangular fluorescence data from [24] for marked human kinetochore proteins, analysing three fluorophore triplets amongst the four fluorescent markers shown schematically in Fig 5. These include fluorophores near the N and C termini of the Ndc80 complex, a fluorophore on Mis12, a protein that binds Ndc80, extending its rigid arm, and CenpC. There are two conditions - standard growth conditions (DMSO) and under nocodazole treatment where microtubules are fully depolymerised; kinetochores are thus in an unattached configuration. We applied an additional quality control step on the data over that employed in [24], requiring that for both sister kinetochores all three fluorophores were observed and that there were at least ten sister pairs in each cell. This was found to reduce the biological variation within an experiment. The same experimental datasets were analysed in [24] with BEDCA; results are similar to the pair-wise results presented here - differences arise because of the extra quality-control filter and the less-informative priors used on the measurement errors in this paper (see supplement S5).

We tested our method on experimental data in three ways: i. confirming zero length is inferred for proteins that are triple labelled (the CenpC–CenpC distance, Example 1), ii. comparing consistency for the posterior length that is common to different triplets (specifically Ndc80C–Ndc80N), iii. comparing with the pair-wise method of [13], see Table 3.

**Table 3.** Single-state analysis of experimental datasets. Examples 1–5 with fluorophore labels, number of kinetochores (*N* ) and number of cells (*C*). Means and standard deviations of the inferred length posteriors are given, comparing the triangle inference presented in this paper with the pair-wise method of [13]. The pair-wise and triangular algorithms were run on the same dataset; the pair-wise results differ slightly to those reported in [24] because of differences in the priors. For the number of kinetochores and cells, see Table S14.2.

| Example 1, CenpC–CenpC–CenpC, $N = 1250$, $C = 32$:<br>(lengths and errors in nm) | | | |
| --- | --- | --- | --- |
|  | CenpC–CenpC | CenpC–CenpC | CenpC–CenpC |
| BEDCA, [13] | $2.9 \pm 1.9$ | $4.4 \pm 2.9$ | $4.7 \pm 3.0$ |
| triangle, uninformative prior | $3.2 \pm 2.1$ | $4.2 \pm 2.7$ | $4.4 \pm 2.7$ |
| Example 2, CenpC–Ndc80C–Ndc80N (DMSO), $N = 72$, $C = 3$:<br>(lengths and errors in nm) | | | |
|  | CenpC–Ndc80C | CenpC–Ndc80N | Ndc80C–Ndc80N |
| BEDCA, [13] | $45.7 \pm 3.2$ | $86.7 \pm 2.5$ | $54.2 \pm 2.9$ |
| triangle, uninformative prior | $43.2 \pm 2.6$ | $86.5 \pm 2.6$ | $49.0 \pm 3.2$ |
| Example 3, CenpC–Ndc80C–Ndc80N (nocodazole), $N = 118$, $C = 5$:<br>(lengths and errors in nm) | | | |
|  | CenpC–Ndc80C | CenpC–Ndc80N | Ndc80C–Ndc80N |
| BEDCA, [13] | $53.1 \pm 8.1$ | $62.1 \pm 3.9$ | $11.6 \pm 7.2$ |
| triangle, uninformative prior | $55.1 \pm 3.5$ | $60.9 \pm 3.6$ | $13.1 \pm 6.7$ |
| Example 4, Nnf1–Ndc80C–Ndc80N (DMSO), $N = 570$, $C = 18$:<br>(lengths and errors in nm) | | | |
|  | Nnf1–Ndc80C | Nnf1–Ndc80N | Ndc80C–Ndc80N |
| BEDCA, [13] | $15.2 \pm 8.3$ | $71.6 \pm 0.8$ | $50.4 \pm 1.0$ |
| triangle, uninformative prior | $30.0 \pm 1.2$ | $71.5 \pm 0.8$ | $47.5 \pm 1.0$ |
| Example 5, Nnf1–Ndc80C–Ndc80N (nocodazole), $N = 238$, $C = 8$:<br>(lengths and errors in nm) | | | |
|  | Nnf1–Ndc80C | Nnf1–Ndc80N | Ndc80C–Ndc80N |
| BEDCA, [13] | $15.3 \pm 9.3$ | $21.0 \pm 11.4$ | $12.7 \pm 7.7$ |
| triangle, uninformative prior | $16.2 \pm 8.4$ | $16.8 \pm 9.5$ | $12.6 \pm 7.4$ |

Firstly, for the triple-labelled CenpC experiment, Example 1 in Table 3, all three lengths of the triangle have posterior marginals that are against the boundary (zero), practically identical to the pair-wise algorithm (see Fig S13.2), giving a resolution of 2–3nm (although this is sample size dependent). Since all three lengths are small here, there is no benefit in using three markers. Secondly, the distance between Ndc80C and Ndc80N is labelled in two triangles with both DMSO and nocodazole treatments, allowing two comparisons. In DMSO, the Ndc80N–Ndc80C agree (Examples 2 and 4, *p* = 30*, see supplement Computation of *p* values for definition of *p*), posteriors are given in Figs 6, 8. Similarly, the Ndc80N–Ndc80C distance agrees in nocodazole, (Examples 3, 5, *p* = 48*), posteriors in Figs 7, S13.3. For the smaller lengths, Ndc80N–Ndc80C in nocodazole, the posterior under the pair-wise correction has a thick tail towards zero, whilst the triangle correction retains a substantial peak, Fig S13.3. This is analogous to the thick tails seen in simulated data for small length inference. Thirdly, the results from the triangle correction are typically consistent with the pair-wise correction method of [13], Table 3. Exceptions are the Nnf1–Ndc80C length (*p* = 1.0*) and the Ndc80C–Ndc80N length (*p* = 1.9*) in the DMSO-treated Nnf1–Ndc80C–Ndc80N, showing a weakly significant difference, see Example 4 in Table 3. Examining the posteriors, Fig 8, reveals that the short Nnf1–Ndc80C length from the pair-wise algorithm is approximately flat between 0nm and 27nm. In contrast, the triangle correction’s length posterior is approximately Gaussian, with substantially smaller posterior variance. The tighter inference of the Ndc80N–Ndc80C distance by the triangle correction is a consequence of length correlations within the inferred triangle. Generating triangles from the pair-wise marginal length posteriors, assuming independence, results in substantial violation of the triangle inequality, Fig 8d. In fact, even the pair-wise posterior means violate the triangle inequality in this case, indicating it is not possible to construct a joint triangle distribution that preserves the pair-wise length marginals (see Triangle length means never violate the triangle inequality for why no triangle distributions exists that has (marginals with) means violating the triangle inequality).

**Fig 6.**
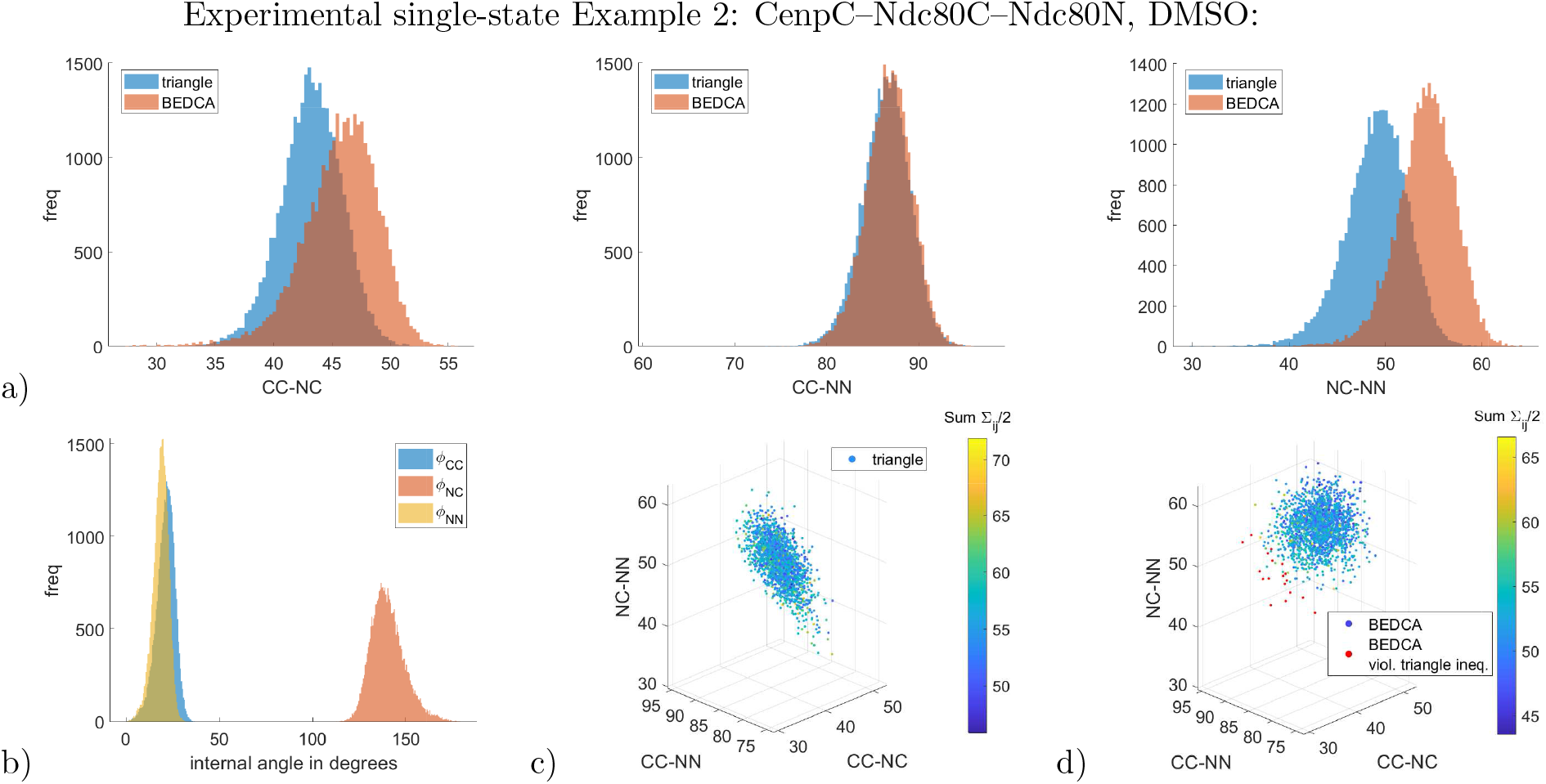
Marginal posteriors of the CenpC–Ndc80C–Ndc80N experiment in DMSO treatment (Example 2 in Table 3). Panels are the same as in Fig 3. Constructing a joint distribution from the three pair-wisely inferred lengths assuming independence yields 1* violations of the triangle inequality (red dots in panel d)).

**Fig 7.**
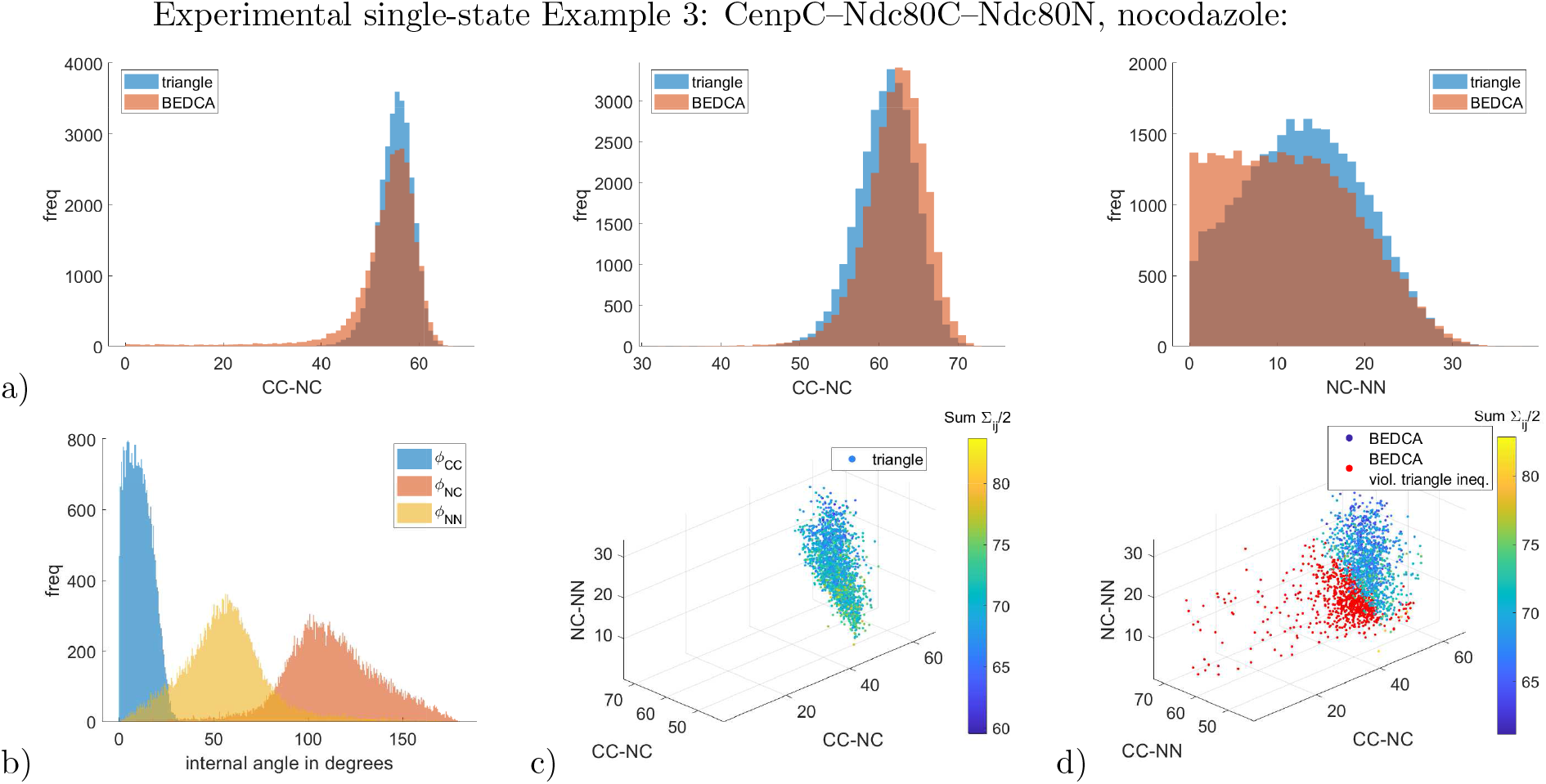
Marginal inferred posteriors of the CenpC–Ndc80C–Ndc80N experiment in nocodazole treatment (Example 3 in Table 3). Panels are the same as in Fig 3. Constructing a joint distribution from the three pair-wisely inferred lengths assuming independence yields 39* violations of the triangle inequality (red dots in panel d)).

**Fig 8.**
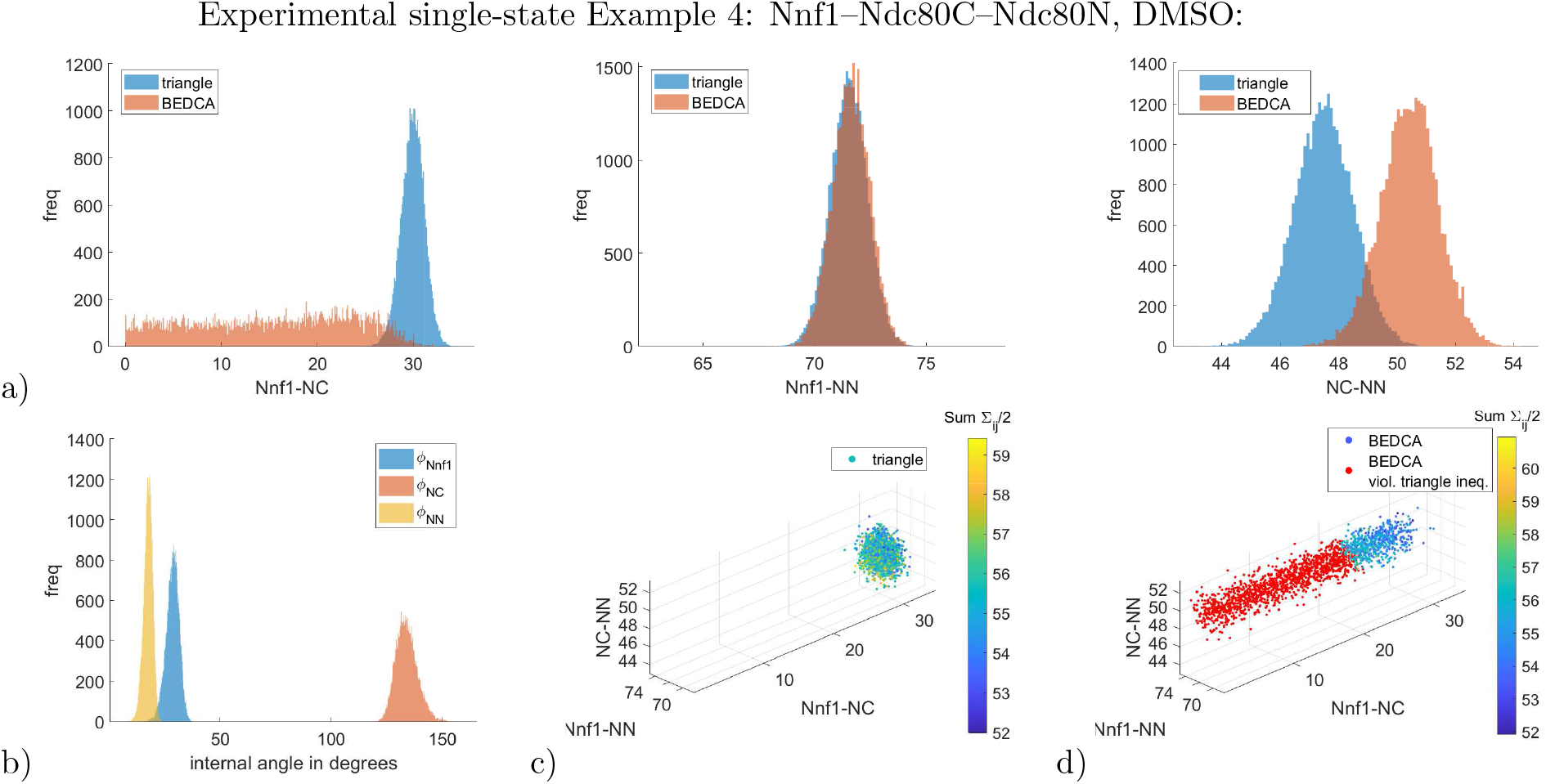
Marginal inferred posteriors of the Nnf1–Ndc80C–Ndc80N experiment in DMSO treatment (Example 4 in Table 3). Panels are the same as in Fig 3. Constructing a joint distribution from the three pair-wisely inferred lengths assuming independence yields 71* violations of the triangle inequality (red dots in panel d)).

The higher confidence of the triangle correction on small lengths reveals that the Nnf1–Ndc80C distance likely increases to twice its length in DMSO compared to nocodazole (Examples 4, 5 in Table 3). This suggests a possible gain of alignment/order, i.e. the ensemble of Ndc80-Nnf1 complexes increase their alignment with each other upon kinetochore attachment to microtubules (see [24] for further discussions).

Triangle correction allows analysis of the measurement errors of each fluorophore individually - see supplementary Table S14.3 for the means and standard deviations of the inferred marginal posteriors. Results are consistent between the examples, if we look at the same fluorophores, molecular structures and treatments. In this data, the CenpC fluorophore (in Examples 2, 3 in Table 3) exhibits the smallest error, having about half the error of some of the other markers. Treatment can also have an effect on the measurement errors.

#### Experimental data for two-state mixture model

Here we analyse heterogeneity of the conformational state of DMSO treated metaphase cells using the data on the CenpC–Ndc80C–Ndc80N and the Nnf1–Ndc80C–Ndc80N triangles that was previously analysed in Examples 2–5, Table 3. As can be seen in Table 3 (and reported previously in [24]), these triangles are distinctly different in attached (DMSO) and unattached (nocodazole) conditions. However, DMSO treated metaphase cells are likely to comprise a mixture of bi-oriented kinetochores (major population, attached) and a minor population of unattached kinetochores that would be in the “jackknifed state”, potentially similar to nocodazole treated kinetochores. The existence of such a population has not been confirmed. Here, we use our triangle mixture algorithm to determine if we can detect such a population.

Firstly, we tested the algorithm’s ability to detect minor populations on real data by creating an artificial mixture using the Nnf1–Ndc80C–Ndc80N fluorophore triplet (Example 1, Table 4). We first split the kinetochores from DMSO treated cells (Example 4, Table 3) into two parts, *N* = 382 and *N* = 188 kinetochores. The first group is analysed using the single-state model to generate priors for the lengths and measurement errors, Eq (8) (and supplement Inference results on subset of Nnf1–Ndc80C–Ndc80N). The second group is pooled with a specified proportion of kinetochores from nocodazole treated cells. The resulting mixed dataset contains a known infused proportion of 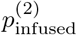 of jackknifed states, in addition to any sub-population in a nocodazole-like state already present. We analysed this mixed dataset with the two-state model, using the remainder of the nocodazole dataset as a pure sample of the jackknifed state *ζ* = 2. The CenpC–Ndc80C–Ndc80N datasets (Examples 2–3, Table 3) were of insufficient size to enable a similar analysis to be performed.

**Table 4.** Two-state experimental examples. Three examples of inference of two states in a mixed state population using the experimental data of Table 3. Inferred posterior means and standard deviations of the triangle lengths of the two states are shown. Kinetochore and cell sample sizes are given in column 10 as: mixed dataset *N*_mix_, informing dataset *N*_inform_, (cell number *C*). Abbreviations are CC for CenpC, NC for Ndc80C and NN for Ndc80N. In Example 1 the DMSO data is infused with the stated proportion 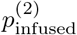 of nocodazole treated kinetochores. In Example 1 the informative prior from Eq (8) is used. In Examples 2 and 3 the DMSO datasets are analysed for a mixture of states, with the nocodazole data used as the informing dataset for the minority state. The priors are uninformative. For number of kinetochores and cells, see Table S14.2.

| Example 1, Nnf1–Ndc80C–Ndc80N, synthetic heterogeneous mixture (DMSO kinetochores infused with jackknifed kinetochores): |  |  |  |  |  |  |  |  |  |
| --- | --- | --- | --- | --- | --- | --- | --- | --- | --- |
| (lengths and errors in nm) | state 1: | | | state 2: | | | $p^{(2)}$ | $p_{\text{infused}}^{(2)}$ | sample sizes |
|  | Nnf1–NC | Nnf1–NN | NC–NN | Nnf1–NC | Nnf1–NN | NC–NN |  |  |  |
| triangle, Eq (8) prior | $29.4 \pm 2.1$ | $71.0 \pm 1.5$ | $45.8 \pm 1.8$ | $16.7 \pm 8.4$ | $16.8 \pm 9.5$ | $13.0 \pm 7.6$ | $(2.20 \pm 2.16) \%$ | 0.00% | 188, 238 (14) |
| triangle, Eq (8) prior | $30.2 \pm 2.0$ | $70.0 \pm 1.6$ | $44.3 \pm 1.9$ | $15.5 \pm 8.3$ | $16.5 \pm 9.8$ | $13.4 \pm 7.7$ | $(4.79 \pm 4.03) \%$ | 10.5% | 210, 216 (14) |
| triangle, Eq (8) prior | $28.8 \pm 2.2$ | $68.4 \pm 1.7$ | $44.3 \pm 2.0$ | $16.2 \pm 8.9$ | $16.5 \pm 9.4$ | $12.7 \pm 7.6$ | $(13.7 \pm 6.9) \%$ | 19.7% | 234, 192 (14) |
| triangle, Eq (8) prior | $29.7 \pm 2.2$ | $68.8 \pm 1.8$ | $43.4 \pm 2.2$ | $15.7 \pm 8.6$ | $15.8 \pm 9.2$ | $13.9 \pm 7.9$ | $(22.6 \pm 7.9) \%$ | 29.9% | 268, 158 (14) |
| triangle, Eq (8) prior | $28.4 \pm 2.3$ | $66.8 \pm 1.9$ | $42.7 \pm 2.3$ | $17.5 \pm 9.1$ | $16.9 \pm 9.9$ | $14.3 \pm 8.2$ | $(41.4 \pm 7.5) \%$ | 40.1% | 314, 112 (14) |
| Example 2, CenpC–Ndc80N–Ndc80C, DMSO treated: |  |  |  |  |  |  |  |  |  |
| (lengths and errors in nm) | state 1: | | | state 2: | | | $p^{(2)}$ | $p_{\text{infused}}^{(2)}$ | sample sizes |
|  | CC–NC | CC–NN | NC–NN | CC–NC | CC–NN | NC–NN |  |  |  |
| triangle, uninformative prior | $42.9 \pm 2.7$ | $86.6 \pm 2.6$ | $49.2 \pm 3.3$ | $55.1 \pm 3.5$ | $61.0 \pm 3.7$ | $13.2 \pm 6.8$ | $(3.67 \pm 3.64) \%$ | 0.00% | 72, 118 (8) |
| Example 3, Nnf1–Ndc80C–Ndc80N, DMSO treated: |  |  |  |  |  |  |  |  |  |
| (lengths and errors in nm) | state 1: | | | state 2: | | | $p^{(2)}$ | $p_{\text{infused}}^{(2)}$ | sample sizes |
|  | Nnf1–NC | Nnf1–NN | NC–NN | Nnf1–NC | Nnf1–NN | NC–NN |  |  |  |
| triangle, uninformative prior | $30.4 \pm 1.2$ | $71.7 \pm 0.8$ | $47.7 \pm 1.0$ | $14.0 \pm 8.0$ | $15.7 \pm 9.3$ | $12.3 \pm 7.4$ | $(3.47 \pm 2.06) \%$ | 0.00% | 570, 238 (26) |

There was good agreement between the infused proportion of nocodazole kinetochores, 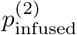 , and the inferred proportion of nocodazole-like states, *p*^(2)^; means and standard deviations are given in Table 4 and marginal posteriors in Fig 9. For infused proportions larger than 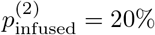 the weight of the marginal posterior of *p*^(2)^ moves away from zero indicating the presence of the minor population. The plot of inferred proportion against infused proportion shows the expected linear relationship, Fig 9b. The lengths and measurement errors are inferred accurately at the lower infusion levels (compared to the single state triangle correction results, subsection Experimental data for single-state model). However, there is a systematic trend in the inferred Nnf1–NN length in state *ζ* = 1 towards lower values, thus becoming more similar to the length estimated for the jackknifed state, *ζ* = 2. At 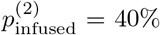 this length becomes significantly different from the state inferred in infusion-free DMSO (Example 3 in Table 4), (*p* = 1.0%). This indicates limitations of this approach; datasets need to be informative enough, e.g. by using sufficiently high sample sizes, or by using sufficiently strong priors on state *ζ* = 1.

**Fig 9.**
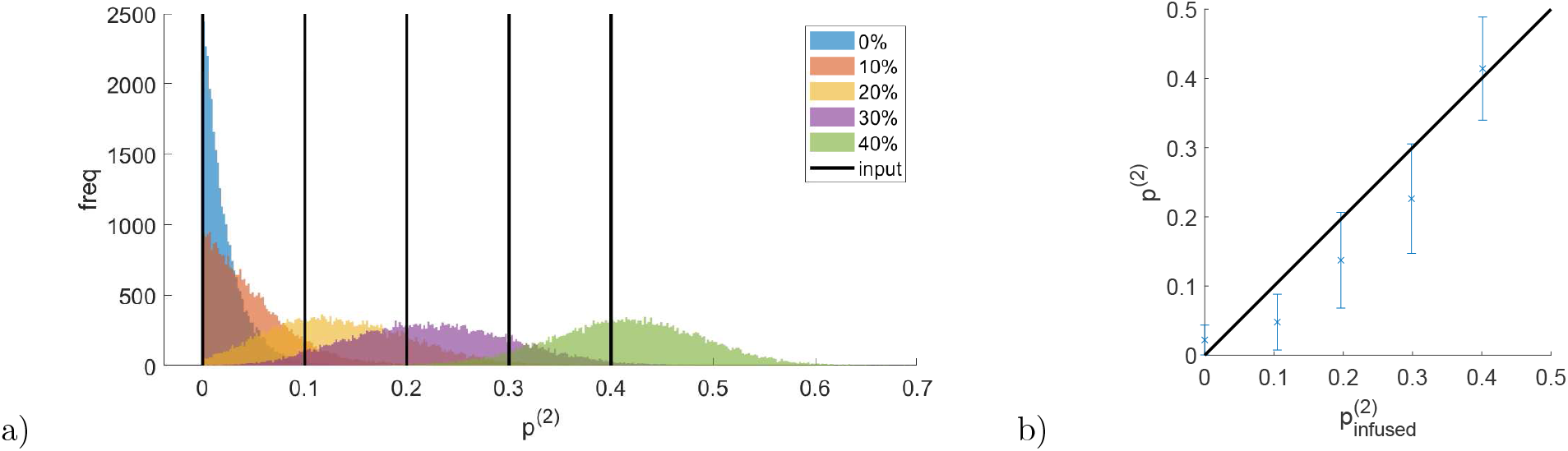
Two-state experimental examples. Two-state mixture inference example with infusion of kinetochores from nocodazole treated cells into a DMSO dataset. (a) the posterior minor state proportion *p*^(2)^ for various infusion levels 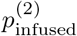. (b) plot of the posterior minor state proportion *p*^(2)^ against the infusion level 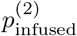. Each point is based on a single realisation of the mixture. The posterior mean is plotted with error bars showing the standard deviation. The black line is 1:1 relationship 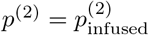.

**Fig 10.**
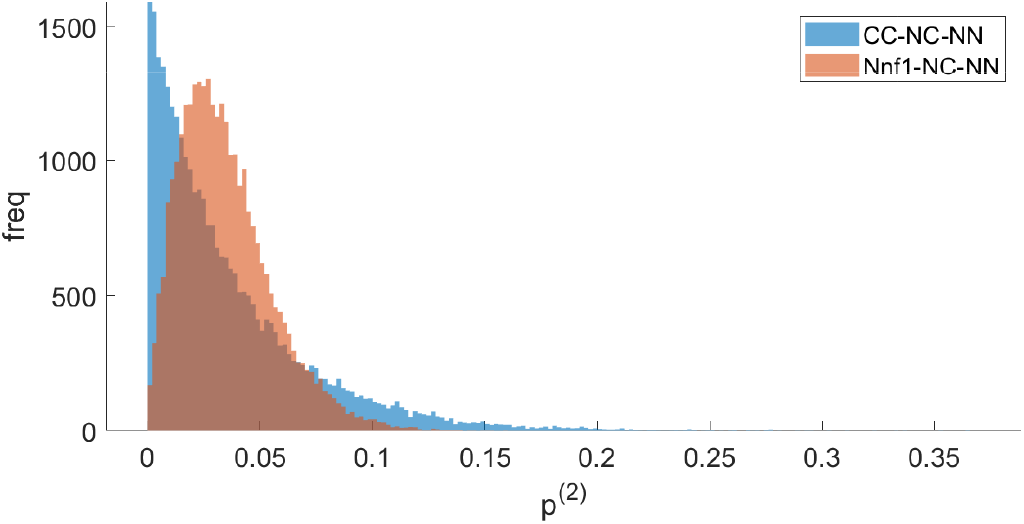
Two-state inference of synthetically mixed experimental datasets. Inferred marginal posteriors of the (minor) state proportion *p*^(2)^ in the two experimental datasets labelled with triplets CenpC-Ndc80N-Ndc80C and Nnf1-Ndc80N-Ndc80C, Examples 2, 3 respectively of Table 4.

Secondly, we analysed the two DMSO triplet datasets with the two-state model, using the nocodazole dataset (assumed pure) to inform the minor population state (*ζ* = 2). We used flat priors (see supplement S1) since the attached states are in the vast majority in the heterogeneous population, Fig 9b. The smaller dataset, Example 2, Table 4 (CenpC–Ndc80C–Ndc80N with *N*_mix_ = 72, *N*_inform_ = 118 kinetochores) has a state proportion *p*^(2)^ posterior with a mode at zero and a tail extending beyond 15%, see Fig 9. Thus, it is inconclusive as to whether a minor population exists. However, for the larger dataset, Example 3, Table 4 (Nnf1–Ndc80C–Ndc80N with *N*_mix_ = 570, *N*_inform_ = 238 kinetochores) the proportion *p*^(2)^ posterior separates away from zero, Fig 9 with inferred proportion *p*^(2)^ = (3.47 ± 2.06) %. Although inconclusive, the CenpC–Ndc80C–Ndc80N proportion is consistent, Table 4. We thus have strong evidence for DMSO treated cells in metaphase being heterogeneous with respect to Ndc80 conformation. A model comparison indicated a probability *p* = (77.9 ± 4.0) % in favour of the multi-state model compared to the single-state model. See supplement Model comparison: Two-state vs single-state model for details.

Although with our method the mixture proportions {*p*^(*ζ*)^}_*ζ*_ can be inferred accurately (verified in simulated data, synthetic mixtures), unfortunately there is insufficient information in our examples to identify the state *ζ*^*n*^ of an individual kinetochore *n*, see supplementary Fig S12.1.

## Conclusion

Here we have presented a Bayesian methodology to infer the 3D polygon geometry of *J* ≥ 2 fluorophores within a macromolecular complex, for markers that localise to distinct regions within the complex and are spot-like when imaged. The algorithm corrects for length and angle bias due to measurement noise, biases that become substantial when polygon sides are of the order of, or less than, the measurement error. Previous correction methods ( [6], [5], [13]) were pair-wise methods. Our new method improves on these in four principal ways:

- The full information in multi-fluorophore images is used to infer the polygon, allowing internal angles between the polygon edges and correlations between edge lengths to be inferred.
- Lengths can be inferred with higher confidence if more than two markers are used. This is particularly beneficial for short lengths (relative to the measurement error), and can overcome inference problems previously reported by [20] (in a single-molecule context). The largest benefits occur when the third marker is located far away and the three markers are approximately co-linear.
- Using three or more fluorophores allows measurement errors to be inferred individually for each marker. This could for example be used for quality control of fluorophores or for quantifying spot expansions (provided the measurement errors remain approximately Gaussian), such expansions potentially being of biological relevance.
- The method is highly flexible. Our algorithm allows for anisotropic measurement errors, typical for 3D imaging, and extends to heterogeneous datasets, inferring both the majority state and the composition of a mixture of multiple states.

Our algorithm models the full system geometry, which gives it great flexibility, allowing for further generalisations: For instance non-Gaussian measurement errors or anisotropic orientations *R*^*n*^ would be easy to implement. Anisotropy of orientations is likely common in 3D imaging; for example cells may be selected with spindle poles approximately within a focal plane - since kinetochores have a preference to lie along the spindle axis when bi-orientated they are then not likely to be orientated isotropically as a population. This flexible approach does however come with longer run-times compared to the pair-wise methods ( [13], [5]). The run-time of the single-state simulated Example 1 (*N* = 400 measurements and 100, 000 Markov chain iterations) was about 2 days. This could be reduced by using state-of-the-art samplers, *e.g.* sequential Monte Carlo, Hamiltonian Monte Carlo, [1, Ch 2,9], a re-parametrisation that captures system structural correlations (e.g. utilises the relation in S6), or parallelised algorithms [4] (potentially of great benefit since the perspectives, {*T*^*n*^}_*n*_, {*R*^*n*^}_*n*_, are updated independently).

The two-state algorithm enabled us to analyse heterogeneities in the architecture of human kinetochores in metaphase, providing direct evidence for a minor population of kinetochores in an unattached conformation. Our method allows conformations of macromolecular complexes to be analysed offering a new technique for study of conformational change in vivo, distinct from conformation (proximity) sensitive readout approaches. These include for example Double Electron-Electron Resonance (DEER; e.g. [16]), Förster Resoncance Energy Transfer (FRET; e.g. [15]) and Bimolecular Fluorescence Complementation (BiFC; e.g. [23]). Our method is a post-processing method for multi-fluorophore image data and does not need a specialised experimental setup or specialised fluorophores as these other methods. Our multi-state algorithm requires a large enough sample size to allow for the statistical analysis (in our examples tens to hundreds of samples) and a discrete number *Z* of distinct (polygon) states. For our examples we needed to provide information on rare state(s) within a heterogeneous dataset which we achieved by using joint inference on two datasets (adding an informing dataset). It is likely that the decomposition of heterogeneous populations can be improved, including identification of individual sample state, by using higher-resolution imaging, e.g. Super-Resolution Structured Illumination Microscopy (SR-SIM, [17]), Super-Resolution Radial Fluctuations (SRRF, [9]) microscopy, or by capturing substantially more photons. On the other hand, our method works at any distance, and while resolution decreases for shorter lengths, on some examples we get confidences as low as 1nm. This range of tens of nanometres is important at the molecular scale. For instance the conformation change of the kinetochore from attached to unattached state involves a reduction of the CenpC–Ndc80N distance from 85nm to 60nm, which is beyond the scope of DEER or FRET. From this perspective our multi-state algorithm complements these experimental methods, as it bridges the gap between their S 10nm range and the length scale of the PSF, where distance corrections become negligible.

## Supporting information

### S1. Uninformative priors

We use uninformative (improper) priors, namely:

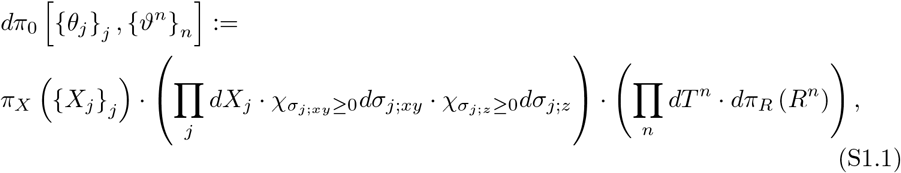

where *χ* is the characteristic function, *π*_*X*_ [{*X*_*j*_}_*j*_] is the prior distribution on the template positions and *dπ*_*R*_ [*R*^*n*^] is the probability measure on the rotation matrices *R*^*n*^ ∈ *SO* (3) as follows.

Uninformative priors are typically chosen to be flat (uniform); however this is not re-parametrisation independent, and there are many choices of polygon properties where a uniform prior can be imposed. Because our interest is in the polygon side lengths, a flat prior *l*_*ij*_ ≔ |*X*_*j*_ − *X*_*i*_| would appear appropriate, giving the (improper) prior ∝ (∏_*i*≠*j*_ *dl*_*ij*_). However, the marginal prior for any length would in fact not be flat for *J* > 2, because of the constraints on the lengths to form a *J*-polygon. Priors with approximately flat marginal densities in the lengths can be constructed for the triangle case (*J* = 3) using the density (see Flat prior on marginals of lengths for a derivation):

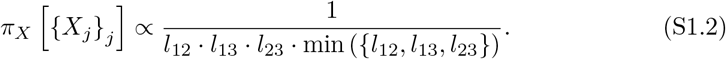

Three priors on the template positions are compared in Fig S1.1, namely (∏_*j*_ *dX*_*j*_), 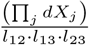 and *π*_*X*_ [{*X*_*j*_}_*j*_] · (∏_*j*_ *dX*_*j*_).

The measure *dπ*_*R*_ on the rotation matrix *R*^*n*^ is defined, such that for an arbitrary vector on the ℝ^3^-unit sphere, 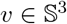, the image *R*^*n*^ · *ν* · *dπ*_*R*_ [*R*^*n*^] is uniform on the surface of 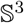 (i.e. the Lebesgue measure on that surface, sin(*ψ*)*dψdφ* in spherical polars).

### S2. Flat prior on marginals of lengths

Here we show, that the distribution given in Eq (S1.2) corresponds to a flat prior on the marginals of the lengths in the limit of unconstrained triangle sizes.

Triangles are specified by the three 3D positional vectors *X*_*j*_ , *j* ∈ {1, 2, 3}, giving the unbounded parameter space ℝ^9^. Consider the stripe subspaces,

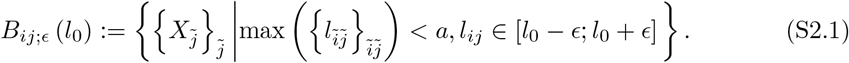

corresponding to length *l*_*ij*_ ≔ |*X*_*j*_ − *X*_*i*_| constrained to a small *ϵ*-neighbourhood of a fixed value *l*_0_ > 0, and all lengths being less than *a >* 0. This stripe region is infinite, because of the translational degrees of freedom, but becomes finite once the translations are fixed. We want to show that for the prior from Eq S1.2 the probability to be in such a domain (and constraining *X*_1_ to a bounded domain) is independent of the value *l*_0_ in the limit of narrow stripes and unbounded triangles:

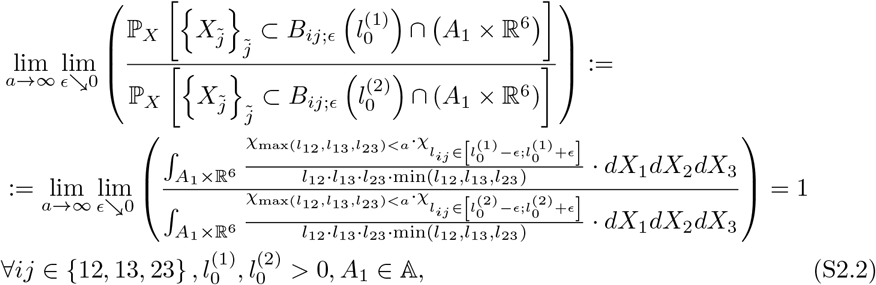

where we used 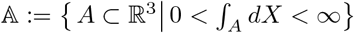 to confine node *X*_1_ to a domain of finite volume.

**Fig S1.1.**
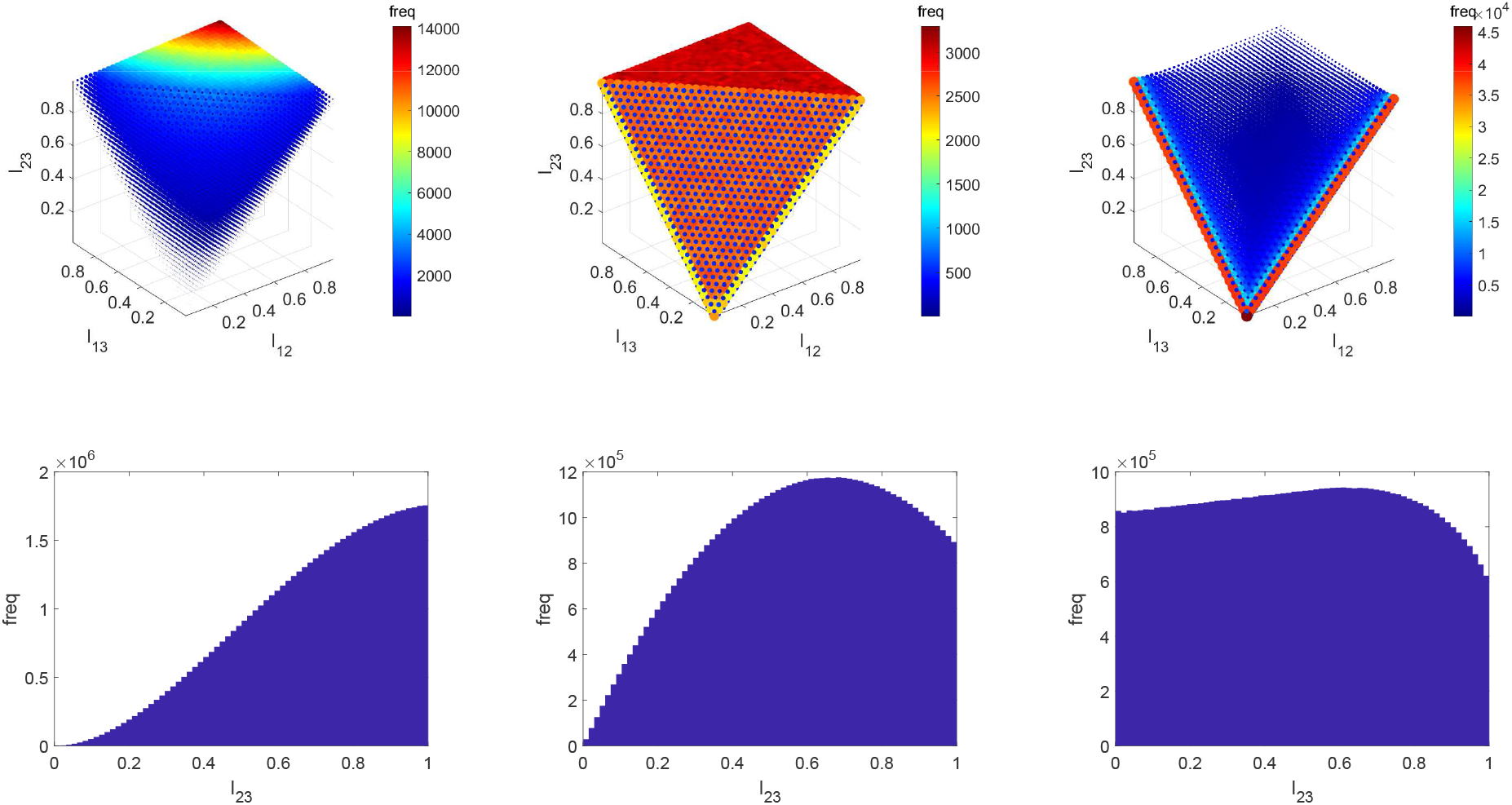
Numerical simulation of the histograms for three possible priors on the triangle template positions. {*X*_*j*_}_*j*_. Marginal densities are shown on the space of the three triangle lengths (top row; frequency is indicated by colour and radius) and the associated marginal distribution of a single length (bottom row). From left to right the prior distributions are 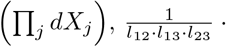 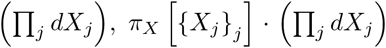. The support is bounded by the triangle inequality, e.g. the front face is the bounding simplex *l*_23_ = *l*_12_ + *l*_13_. A cut-off for each of the lengths was used, forcing them to be within [0, 1]. The symmetry of the prior with respect to re-labelling markers *j* was used when sampling.

We first show, that

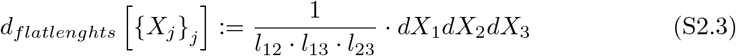

gives a flat prior on the lengths space (within the boundaries of the triangle inequality), i.e.:

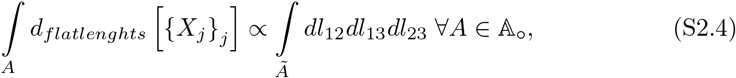

with 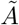 the set of all lengths for which {*X*_*j*_ }_*j*_ ∈ *A* and

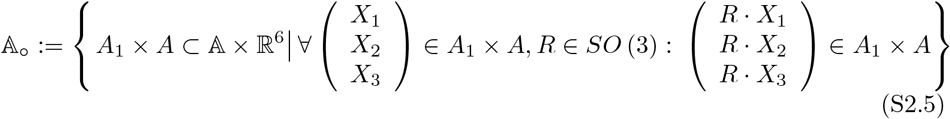

the set of all rotation symmetric positions with 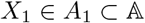. It is well-known that - for two positions *X*_1_, *X*_2_ ∈ ℝ^3^ - we can switch from Cartesian to spherical coordinates via:

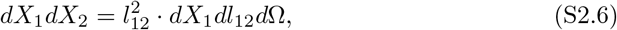

where *d*Ω = sin (*ψ*) · *dφdψ* is the differential of the solid angle (for azimuth and inclination angles *φ*, *ψ*). On the other hand, we know from the transition from Cartesian to cylinder coordinates (with the *X*_1_–*X*_2_-edge as the axis of the cylinder), that *dX*_3_ = *h*_3_ · *dh*_3_*dadθ*, where *h*_3_ ≥ 0 is the distance of *X*_3_ from the cylinder axis, *a* ∈ ℝ is the coordinate along the cylinder axis and *θ* ∈ [0, 2*π*] is the rotation angle of *X*_3_ around the cylinder axis (relative to some fixed plane containing that axis). Expressing *l*_13_, *l*_23_ through the cylinder coordinates *a*, *h*_3_ as well as *l*_12_ we have:

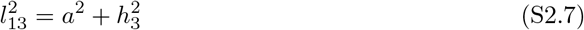

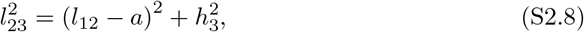

yielding:

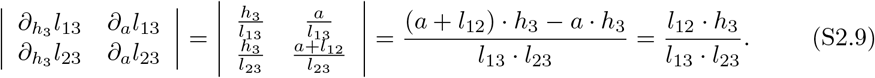

Combining this with our results for *dX*_1_*dX*_2_ and *dX*_3_ we get:

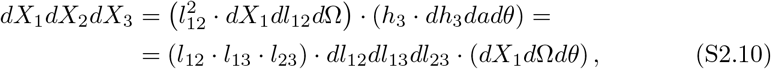

where the term in the last bracket just gives a constant, thus proving Eq (S2.4).

Without loss of generality, we show Eq (S2.2) for *ij* = 23, i.e. the marginal over *l*_23_. Note, that the additional factor min (*l*_12_, *l*_13_, *l*_23_)^−1^ in Eq (S1.2) versus *d*_*flatlengths*_ merely comes from the lengths space only being occupied within the boundaries of the triangle inequality (other points are not contained in 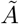 in Eq (S2.4)). To see how the extra factor fixes this, we assume an upper cut-off for all lengths *a* ≫ *l*_0_ > 0 and divide the marginal integral into different parts depending on which length is shortest:

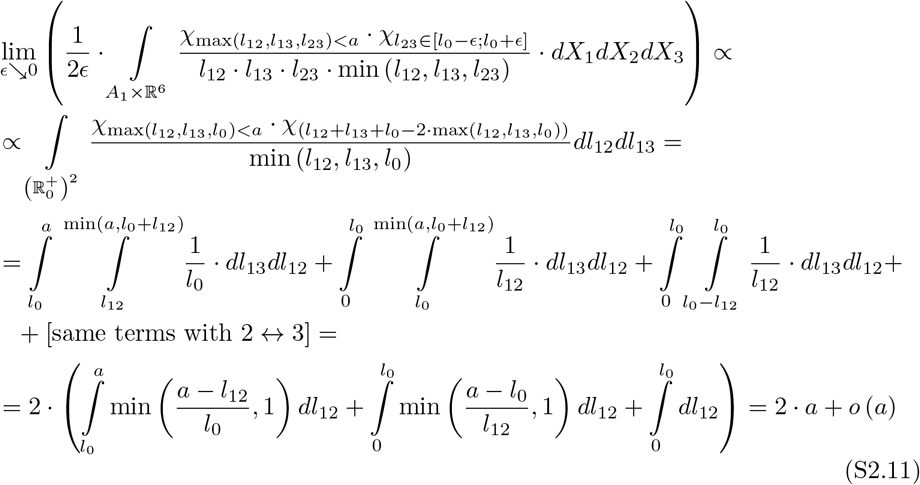

where the terms in the third row correspond to *l*_0_ ≤ *l*_12_ ≤ *l*_13_, *l*_12_ ≤ *l*_0_ ≤ *l*_13_ and *l*_12_ ≤ *l*_13_ ≤ *l*_0_ in this order (and then similarly with 2 and 3 swapped). For *a* → ∞ (i.e. no cut-off of lengths) the *l*_0_-dependent terms *o* (*a*) become negligible, thus proofing our claim from Eq (S2.2).

Note firstly, this is only an asymptotic argument and for finite cut-offs there are slight variations (see Fig S1.1 for a numerical study of the effect of a finite support, *a* < ∞).

Second, the prior on the triangle lengths used here is not unique under the requirement of flat marginals (e.g. a homogeneous weight on the line *l*_12_ = *l*_13_ = *l*_23_ would also satisfy this condition). We also want to highlight that the chosen prior on the lengths is scale invariant, i.e. for a scale parameter *α* ∈ ℝ / {0} the ratio 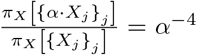 is only a function of *α*, hence does not depend on the triangle shape.

### S3. Markov Chain Monte Carlo samplers for parameter inference

Here we give the details of the Markov Chain Monte Carlo approach taken to sample from the posterior, firstly for the single-state model in Eq (9) and secondly for the multi-state model.

#### S3.1. Samplers of single-state model

We sequentially update the various parameters, {*X*_*j*_}_*j*_ , {*τ*_*j*_}_*j*_ , {*T*^*n*^}_*n*_ , {*R*^*n*^}_*n*_, using the following samplers:

##### Random walk samplers

Random walk samplers are used for the template positions and perspective parameters, implemented sequentially for each of the parameters *X*_*j*_, *T*^*n*^ and *R*^*n*^. We use block updates, blocking together the three coordinates in each of the vectors *X*_*j*_, and block together the six parameters in *T*^*n*^ and *R*^*n*^. For the template positions *X*_*j*_ and translations *T*^*n*^ the proposals are uniform balls around the current position. For the rotations *R*^*n*^ the proposal is implemented as follows:

- choose a reference vector 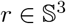 uniformly from the unit-sphere
- draw a sphere location 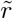 displaced from *r* by choosing it uniformly within a spherical cap around *r* (the radius of the cap is given by the step-size (adjusted during burnin)).
- determine a rotation perturbation 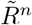 as the “geodesic” rotation that maps *r* onto 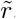, i.e. the rotation with invariant axis 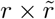 around the angle arccos 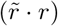
- the proposed rotation is the combined 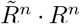, where *R*^*n*^ is the current rotation.

During burnin the step-sizes of all random walk samplers are adjusted with a target rejection rate of *ρ*_*μ*_ ≔ 77.5% (motivated by [26]). The step-size adjustment procedure (given a current step-size of *s* > 0, current adjustment factor of *a* > 1, current deviation *δρ* ∈ ℝ from target acceptance rate *ρ*_*μ*_ and a tolerance of *ρ*_*σ*_ ≔ 7.5%):

- compute the rejection rate (of the random walk update of the respective sampler) *ρ*′ since the last step-size evaluation
- compute the new deviation 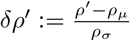
- square-root the adjustment factor *a*, if the deviation *δρ* has the opposite sign to that of the previous update, i.e. if sign (*δρ*) ≠ (*δρ*′)
- get the new step-size as *s*′ ≔ *s* · *a*^−*δρ*^′
- replace *s*, *δρ* for their updated versions *s*′, *δρ*′

We initialise with *a* = 2 and the above five-step procedure was repeated every 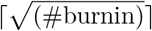 Markov chain iterations through burnin, and initial step sizes are chosen heuristically in the range of 10^−2^ and 10 (in nm for lengths, measurement errors).

##### Gibbs sampler for the translations *T*^*n*^

The translations *T*^*n*^ are also updated using a Gibbs sampler. Here we sample from the target:

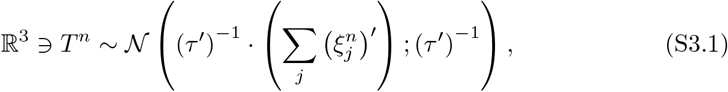

where:

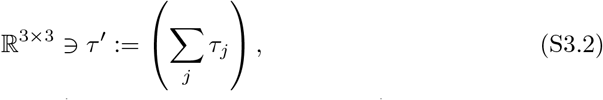

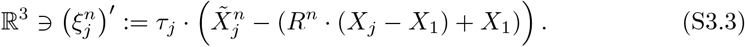

##### Gibbs sampler for the measurement errors *τ*_*j*_

A Gibbs sampler is used for the precisions 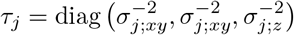 of the measurement errors of the three markers. Note that the two components *xy*, *z* as well as the three markers *j* are independent from each other, so sequential and joint updates coincide. We have a Γ-distribution for each of the conditional distributions:

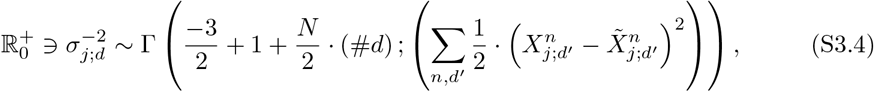

where *d* ∈ {*xy, z*} and number of dimensions (#*d*) = 2, 1, respectively.

##### Initialisation and convergence monitoring

Unless otherwise stated, the variables of all chains are initialised randomly as follows (independent for each *j*, *n*):

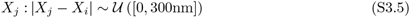

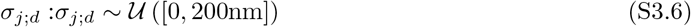

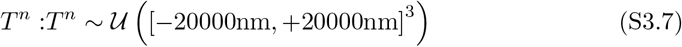

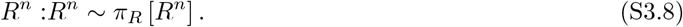

These are over-dispersed compared to the anticipated width of the posterior distribution (confirmed after the run). The total number of iterations per Markov chain for each example is reported in Table S3.1; sub-sampling (equally spaced) was used to give a final sample size of no more than 10000 samples.

A multi-chain convergence diagnostic was used (5 independent chains), assessing convergence by computing the Gelman-Rubin statistic 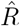 (uncorrected) separately on parameters of interest, specifically |*X*_*j*_ − *X*_*i*_|, *σ*_*j*;*d*_ for *i, j* ∈ {1, 2, 3}, *d* ∈ {*xy, z*}, (see [12, Ch 11.6]). We used a threshold of 1.1; if converged, the five chains were then pooled to reconstruct the posterior.

The computation time on an ordinary desktop computer is about 1 day for datasets with *N* = 200 samples (our Matlab implementation may be further improved for speed).

**Table S3.1.** Number of Markov chain iterations in the examples presented in the main part. In all examples the first 40% of the iterations is taken as burn-in, and the remaining 60% constitute the posterior samples.

| Single-state simulated examples (see Table 1 for results): |  |  |  |  |  |  |
| --- | --- | --- | --- | --- | --- | --- |
| Example | 1 | 2 | 3 | 4 | 5 | 6 |
| number of MCMC iterations | 100000 | 100000 | 500000 | 500000 | 500000 | 500000 |

| Two-state simulated examples (see Table 2 for results): |  |  |  |  |  |  |
| --- | --- | --- | --- | --- | --- | --- |
| Example | 1 | 2 | 3 | 4 | 5 | 6 |
| number of MCMC iterations | 50000 | 50000 | 50000 | 50000 | 50000 | 50000 |

| Single-state experimental examples (see Table 3 for results): |  |  |  |  |  |
| --- | --- | --- | --- | --- | --- |
| Example | 1 | 2 | 3 | 4 | 5 |
| number of MCMC iterations | 100000 | 100000 | 100000 | 100000 | 500000 |

| Two-state experimental examples (see Table 4 for results): |  |  |  |
| --- | --- | --- | --- |
| Example | 1 (all) | 2 | 3 |
| number of MCMC iterations | 200000 | 200000 | 200000 |

#### S3.2. Samplers of multi-state model

All parameters already present in the single-state model from subsection Polygon parametrisation and inference can be inferred using the same updates, if confining the observed data points to the subset in the currently updated state *ζ*^*n*^ = *ζ*. The new variables {*ζ*^*n*^}_*n*_ and {*p*^(*ζ*)^}_*ζ*_ are updated sequentially with random walk and Gibbs samplers, respectively.

##### Random walk sampler for state-affiliations {*ζ*^*n*^}_*n*_

The hidden state-affiliations {*ζ*^*n*^}_*n*_ are sampled with a random walk proposal, equiprobable on all states except the current one (assigned zero probability). To achieve a higher acceptance rate for a new state-affiliation proposal, the translation *T*^*n*^ for observation *n* are altered such that the centre of mass of the triangle (with equal weights for all markers *j*) of the proposed true positions 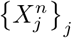 remain unchanged.

##### Gibbs sampler for state proportions 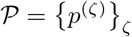

The state proportions 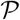 are sampled via a Gibbs sampler from a Dirichlet distribution:

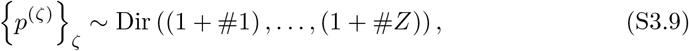

where #*ζ*′ ≔ |{*n* ∈ {1, … , *N*}| *ζ*^*n*^ = *ζ*′}| are the number of measurements in the respective states. The Dirichlet probability density for the state proportions {*p*^(*ζ*)^}_*ζ*_

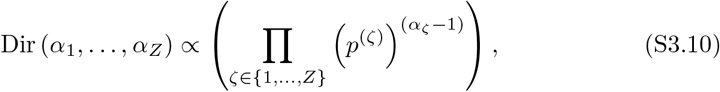

with the state proportions all constrained within [0; 1] and summing to one.

##### Initialisation and convergence monitoring

The parameters were randomly initialised (independent for each *j*, *n*; for Eq (8) priors):

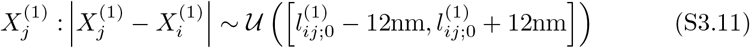

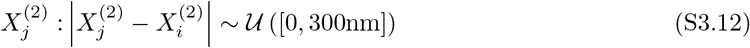

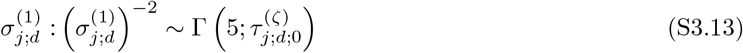

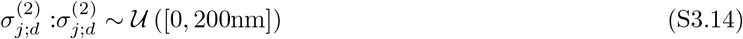

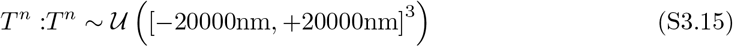

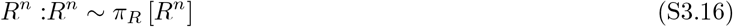

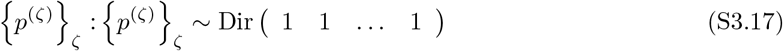

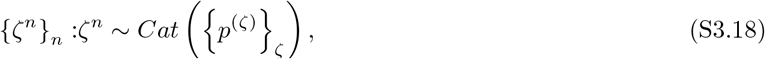

where *Cat* ({*p*^(*ζ*)^}_*ζ*_) is the multinoulli distribution, state *ζ* being drawn with probability *p*^(*ζ*)^. The values 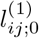 and 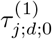 are identical to the priors in Eq (8) and given for each example individually in the main text. These are chosen to be overdispersed compared to the expected posteriors (and confirmed aposteriori), except for the priors for the lengths and measurement errors of the first state. For the effect of the latter on the posterior, see supplement S9.

Convergence is monitored as described above for the single-state model, where additionally each of the *p*^(*ζ*)^ has to stay below the Gelman-Rubin threshold of 1.1. The number of Markov chain iterations for the examples is given in Table S3.1.

### S4. Simulated data

To test the algorithm we simulated data based on the model described in section Materials and methods. For the simulations the template positions 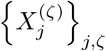, the measurement errors 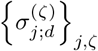 and the state proportions {*p*^(*ζ*)^}_*ζ*_ are fixed as Tables 1, 2. The translations {*T*^*n*^}_*n*_ are independently sampled from Gaussians:

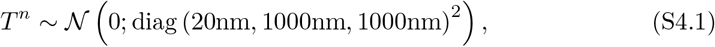

and the rotations are sampled from converged chains of a Markov Chain Monte Carlo algorithm using the same random walk update as described in subsection Samplers of single-state model.

### S5. Implementation of pair-wise correction methods

We compared our method to existing length inference techniques between two markers, namely by Churchman et al from [5] and BEDCA, [13]. Here we describe our implementation of these methods: Churchman et al. [Isotropic measurement errors only]

#### Maximum likelihood estimate (MLE)

we used the in-built Matlab functions `simulannealbnd` then `fminsearch` on the 3D likelihood as specified in Eq (6) in [5]. For error estimates we calculated the Hessian of the likelihood and used its negative for the inverse covariance matrix of the Gaussian approximation.

#### Mean and standard deviation of the full posterior

we used the same likelihood function and assumed flat priors on the lengths |*X*_*j*_ − *X*_*i*_| and measurement errors *σ*_*ij*_. The marginal distributions were computed using numerical integration on a rectangular grid.

##### BEDCA

This is the algorithm used in [13] and [24]; however we used different priors analogous to the ones used in the triangle method (see Eq (S1.1)). Specifically, measurement errors have flat priors on the positive real axis (*dσ*_*j*;*d*_) and lengths have almost flat priors on the positive real axis 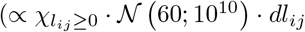 for length *l*_*ij*_ = |*X*_*j*_ − *X*_*i*_|). Note, the latter gives (approximately) flat priors on the lengths (i.e. the length marginals for triangles) but if the three lengths are used to form a triangle under an independence assumption (rejecting those combinations that violate the triangle inequality), the prior would lack the min ({*l*_12_, *l*_13_, *l*_23_}) term from Eq (S1.2) (see subsection Flat prior on marginals of lengths for details).

### S6. Small lengths are increasingly difficult to infer

Here we give an argument, why for a given measurement error, shorter true lengths exhibit a much larger inference error than longer lengths. This phenomenon has been reported in [20] for the length correction based on [5] before.

There are two stochastically independent processes contributing to the distribution of the vectors between two markers, *X*_*j*_ − *X*_*i*_. First, the isotropic distribution of the observed vectors in space (due to the rotations *R*^*n*^) and second, the measurement error (a Gaussian measurement error assumption is not required). Computing the second moment of the distance we obtain:

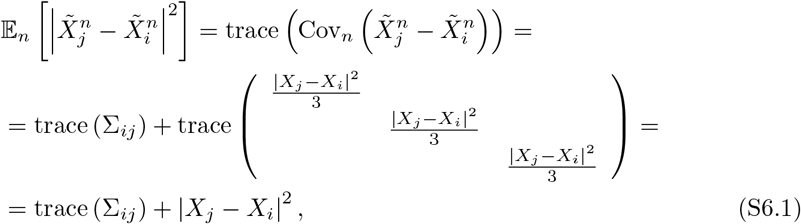

where the expectation and covariance are taken with respect to the marker specific random variables (parametrised with *n*), i.e. measurement errors 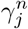, rotations *R*^*n*^ and translations *T*^*n*^; the template positions *X*_*j*_ and measurement errors *σ*_*j*;*d*_ are fixed. In the second line Σ_*ij*_ is the covariance matrix of the measurement error of the displacement (in the setting described in section Materials and methods this would be 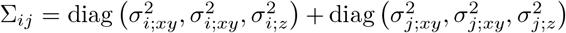 and the matrix in the last term is the covariance of the Lebesgue measure on a sphere of radius |*X*_*j*_ − *X*_*i*_|. Expression (S6.1) is the 3D analogue to Eq (4) in [6]. To quantify how strongly |*X*_*j*_ − *X*_*i*_| depends on the measurement error, we take the derivative of (S6.1):

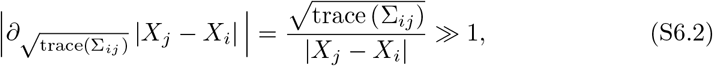

when the true length is much smaller than the measurement error. Thus, small changes in the inferred measurement error lead to very large changes in the inferred true length, with increasing effect for small lengths. See Fig S6.1 for a graphical depiction.

### S7. Computation of *p* values

For comparisons of posteriors, we define the overlap probability

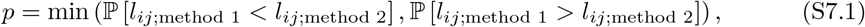

for length *l*_*ij*_ inferred by methods 1, 2. If both methods had identical posteriors, we have *p* = 50%.

### S8. Triangle length means never violate the triangle inequality

For any (proper) probability distribution on the triangle side lengths, *l*_*ij*_ ≥ 0, *i, j* ∈ {1, 2, 3}, the length means necessarily satisfy the triangle inequality (provided they are finite). To prove this, we use Jensen’s inequality ( [3, Thm 3.1.3]) in combination with convexity of the max function to write:

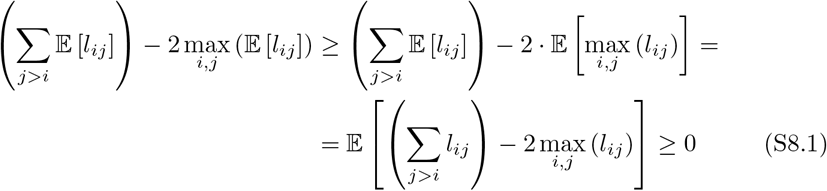

**Fig S6.1.**
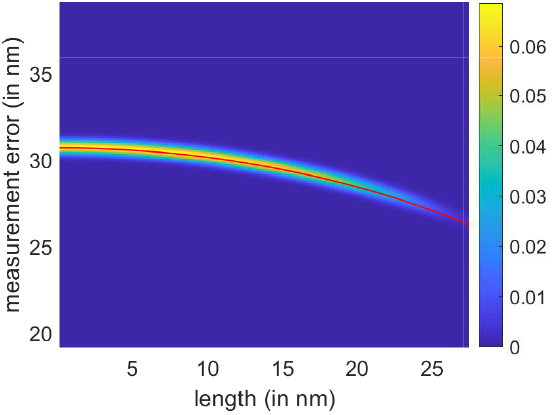
Posterior distribution for measurement error and length for 3D simulated data. Posterior based on *N* = 2000 samples simulated with an input (true) length of 15nm and input (true) measurement error of 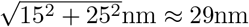. Plotted is the likelihood function, Eq (6) from [5]. Probability density is colour coded as key. The red line is Eq (S6.1) with the second moment (left hand side) estimated from the data.

### S9. Effect of priors in two-state model

For the simulated two-state model and the experimental two-state model with infused nocodazole cells we used informative priors on the *ζ* = 1 triangle state, specifically a box-prior on the side lengths and Gamma-distributed prior on the precisions of the measurement errors (see Eq (8)). Here we analyse the impact of this prior relative to a flat prior on the posteriors of the two-state experimental Example 1 in Table 4 for the cases 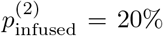 and 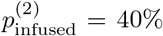; results are summarised in Table S9.1. For 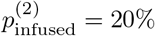 results are very similar (there is a slightly higher confidence for the stronger state-1-prior). For 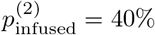 the data is less informative, as there are fewer kinetochores *n* in the first state *ζ* = 1; thus the stronger prior has a greater impact, although results are still consistent.

### S10. Inference results on subset of Nnf1–Ndc80C–Ndc80N

In subsection Experimental data for two-state mixture model we split the Nnf1–Ndc80C–Ndc80N (DMSO-treated) dataset of subsection Experimental data for single-state model into two, one set was used to determine a prior on state *ζ* = 1 for the second set that was used in the two-state model inference. The entire dataset is composed of two independent experiments, with the chronologically determined split occurring within the second experiment, giving subsets of *N* = 382 (in 12 cells; for the prior) and *N* = 188 kinetochores (in 6 cells; for the mixture analysis). The results of the single-state model on the first subset are shown in Table S10.1, rounding them gives the following parameters for an informative prior for state *ζ* = 1 in the two-state model (see Eq (8) for notation):

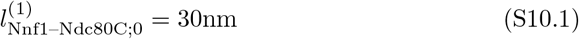

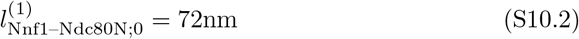

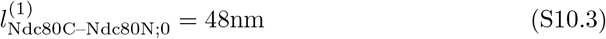

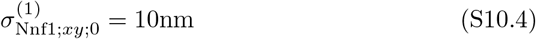

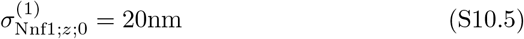

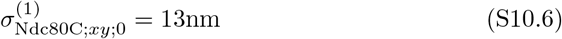

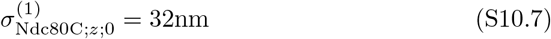

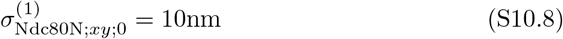

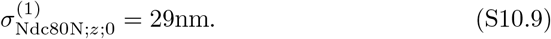

**Table S9.1.** Two-state experimental examples. Comparison of the effect of the “Eq (8) prior” for the two-state model to the flat priors (like for the single-state model) on the inferred lengths, measurement errors and state proportions.

| (lengths and errors in nm) | Nnf1–Ndc80C | Nnf1–Ndc80N | Ndc80C–Ndc80N | $\sigma_{\text{Nnf1};xy}$ | $\sigma_{\text{Nnf1};z}$ | $\sigma_{\text{Ndc80C};xy}$ | $\sigma_{\text{Ndc80C};z}$ | $\sigma_{\text{Ndc80N};xy}$ | $\sigma_{\text{Ndc80N};z}$ | $p^{(\zeta)}$ |
| --- | --- | --- | --- | --- | --- | --- | --- | --- | --- | --- |
| Two-state experimental Example 3, $p_{\text{infused}}^{(2)} = 19.7\%$: | | | | | | | | | | |
| state $\zeta = 1$: | | | | | | | | | | |
| triangle, uninformative prior | $28.5 \pm 2.5$ | $68.2 \pm 1.8$ | $44.5 \pm 2.1$ | $11.0 \pm 2.0$ | $29.8 \pm 3.9$ | $12.3 \pm 1.5$ | $27.1 \pm 2.9$ | $9.8 \pm 2.2$ | $27.2 \pm 3.7$ | $(86.9 \pm 7.5) \%$ |
| triangle, Eq (8) prior | $28.8 \pm 2.2$ | $68.4 \pm 1.7$ | $44.3 \pm 2.0$ | $10.5 \pm 1.5$ | $26.9 \pm 3.8$ | $12.3 \pm 1.3$ | $28.2 \pm 2.4$ | $10.0 \pm 1.5$ | $27.6 \pm 3.0$ | $(86.3 \pm 6.9) \%$ |
| state $\zeta = 2$: | | | | | | | | | | |
| triangle, uninformative prior | $16.3 \pm 9.0$ | $16.4 \pm 9.5$ | $12.7 \pm 7.7$ | $21.1 \pm 2.8$ | $34.0 \pm 3.2$ | $14.5 \pm 3.0$ | $28.1 \pm 2.9$ | $24.5 \pm 1.9$ | $29.1 \pm 2.9$ | $(13.1 \pm 7.5) \%$ |
| triangle, Eq (8) prior | $16.2 \pm 8.9$ | $16.5 \pm 9.4$ | $12.7 \pm 7.6$ | $21.2 \pm 2.7$ | $34.6 \pm 3.2$ | $14.5 \pm 2.9$ | $28.1 \pm 2.9$ | $24.6 \pm 1.9$ | $28.9 \pm 2.9$ | $(13.7 \pm 6.9) \%$ |
| Two-state experimental Example 3, $p_{\text{infused}}^{(2)} = 40.1\%$: | | | | | | | | | | |
| state $\zeta = 1$: | | | | | | | | | | |
| triangle, uninformative prior | $28.1 \pm 2.7$ | $67.1 \pm 2.0$ | $44.5 \pm 2.8$ | $9.7 \pm 2.4$ | $14.8 \pm 8.6$ | $10.2 \pm 2.0$ | $26.5 \pm 3.5$ | $5.9 \pm 3.6$ | $25.2 \pm 6.2$ | $(48.1 \pm 8.8) \%$ |
| triangle, Eq (8) prior | $28.4 \pm 2.3$ | $66.8 \pm 1.9$ | $42.7 \pm 2.3$ | $9.7 \pm 1.5$ | $20.3 \pm 3.8$ | $11.8 \pm 1.3$ | $28.6 \pm 2.6$ | $9.5 \pm 1.5$ | $27.2 \pm 3.7$ | $(58.6 \pm 7.5) \%$ |
| state $\zeta = 2$: | | | | | | | | | | |
| triangle, uninformative prior | $17.8 \pm 9.5$ | $19.0 \pm 10.7$ | $13.5 \pm 8.1$ | $22.0 \pm 3.2$ | $38.2 \pm 3.2$ | $11.8 \pm 3.5$ | $27.3 \pm 3.0$ | $26.8 \pm 1.9$ | $31.6 \pm 3.2$ | $(51.9 \pm 8.8) \%$ |
| triangle, Eq (8) prior | $17.5 \pm 9.1$ | $16.9 \pm 9.9$ | $14.3 \pm 8.2$ | $22.0 \pm 2.7$ | $38.7 \pm 3.2$ | $12.8 \pm 3.6$ | $26.5 \pm 3.4$ | $26.2 \pm 2.0$ | $30.8 \pm 3.4$ | $(41.4 \pm 7.5) \%$ |

**Table S10.1.**
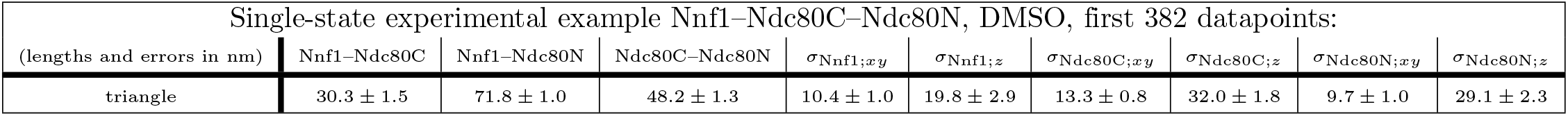
Subset of Nnf1–Ndc80N–Ndc80N, DMSO, data presented in Example 4 of Table 3.

### S11. Model comparison: Two-state vs single-state model

In subsection Experimental data for two-state mixture model we analysed metaphase kinetochores in DMSO with the two-state model (Examples 2, 3 in Table 4). Here we show, how the model comparison with the single-state model was carried out: Apart from the state proportions {*p*^(*ζ*)^}_*ζ*_, the single-state model is a nested sub-model of the two-state model (i.e. it is absolutely continuous) with all state affiliations {*ζ*^*n*^}_*n*_ equal to the same state *ζ*. We can therefore compute the Bayes factor of these two models based on our MCMC samples of the two-state model alone. Let *φ*_*ts*_ and *φ*_*ss*_ denote the model parameters of the two- or single-state models, respectively, except for the state affiliations {*ζ*^*n*^}_*n*_. Their corresponding parameter spaces are denoted Φ_*ts*_, Φ_*ss*_). The Bayes factor is then given by:

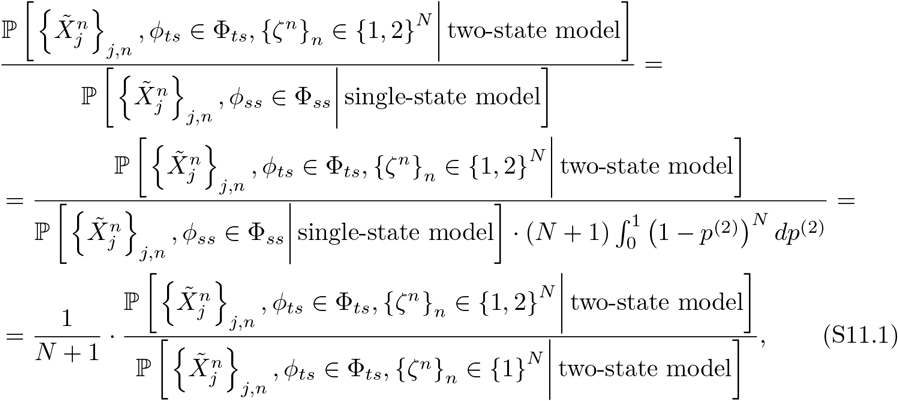

where the last identity comes from the fact that when we confine the multi-state model to the case where all observations are in one state, {*ζ*^*n*^}_*n*_ ∈ {1}^*N*^ , the likelihood times the prior is the same as its single-state counterpart, apart from the extra factor from the Dirichlet distribution, 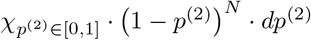 (note we have the same priors on lengths and measurement errors in both models). The right-hand ratio in Eq (S11.1) can be estimated from our MCMC of the two-state model by counting how often the chain is in the pure state 1. We abbreviate this ratio by *ω*. Our discussion so far has assumed a flat prior on the state proportion for the two-state model, 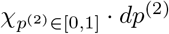. For analysis of a heterogeneous population with a minor sub-population, a flat prior on the smaller interval [0, *α*] for *α* ∈ [0, 1] appears more realistic. In this case the Bayes factor in Eq (S11.1) changes to: 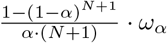.

From the Bayes factor we immediately get the probability of the two-state model (assuming apriori equiprobable models):

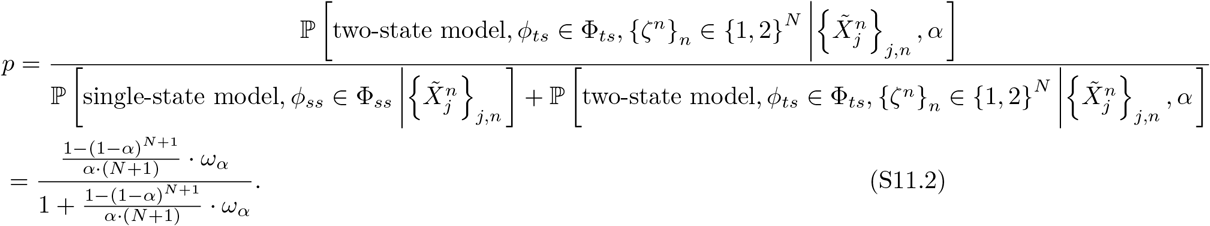

This is plotted as a function of *α* in Fig S11.1 (for Example 3 in Table 4). The *p* value in the main text is computed for *α* = 20%.

### S12. Distribution of the state affiliations {*ζ*^*n*^}_*n*_

In the two-state model we not individual measurement only infer the state proportions {*p*^*ζ*^}_*ζ*_ but also the state affiliations {*ζ*^*n*^}_*n*_ for each individual measurement *n* ∈ {1, … , *N*}. However, the latter do not show a clear bimodal distribution (as a histogram of the mean state affiliation for each measurement *n*) that would allow a clear assignment of individual measurements to a state, see Fig S12.1.

**Fig S11.1.**
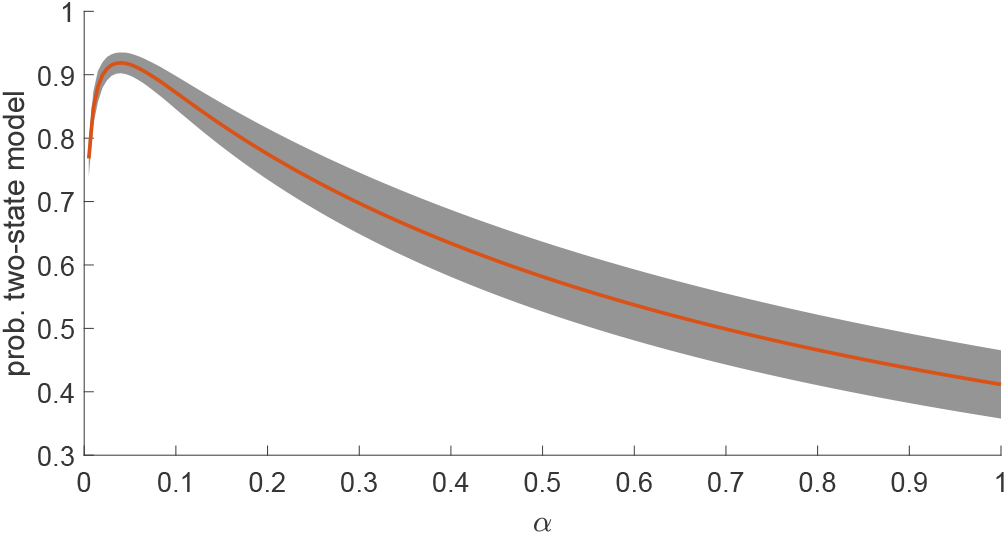
Model comparison between two-state and single-state model for Example 3 in Table 4. Dependence of the probability of the two-state model on the prior parameter *α*, as given in Eq (S11.2). The orange line denotes the mean over the five independent runs, while the shaded area is the ±1*σ* range for each *α*.

**Fig S12.1.**
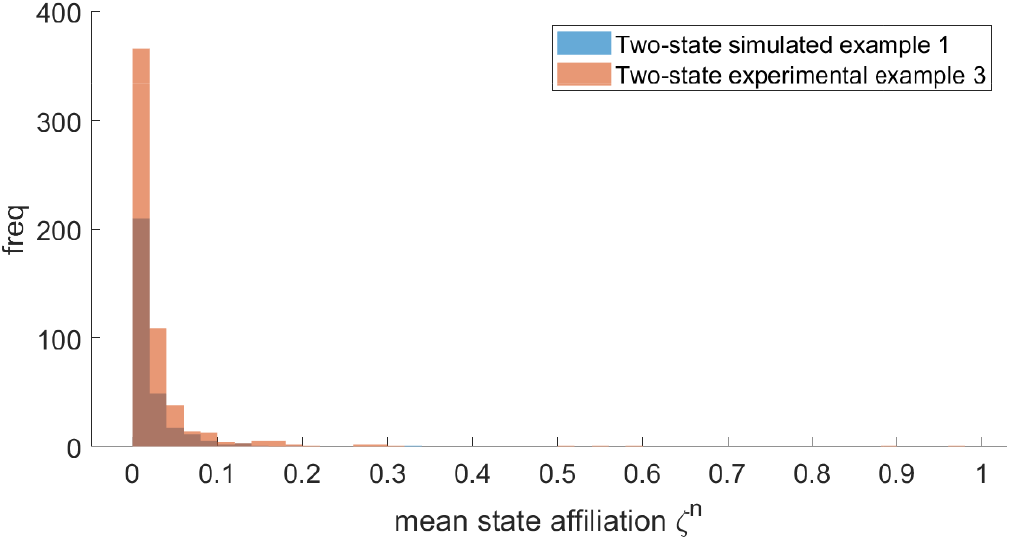
Mean state affiliations {*ζ*^*n*^}_*n*_ for the two-state Examples 1 in Table 2 and 3 in Table 4. Depicted are the means of the state affiliations (over all Markov chain samples) for each of the *N*_mix_ measurements.

### S13. Additional supplementary images

**Fig S13.1.**
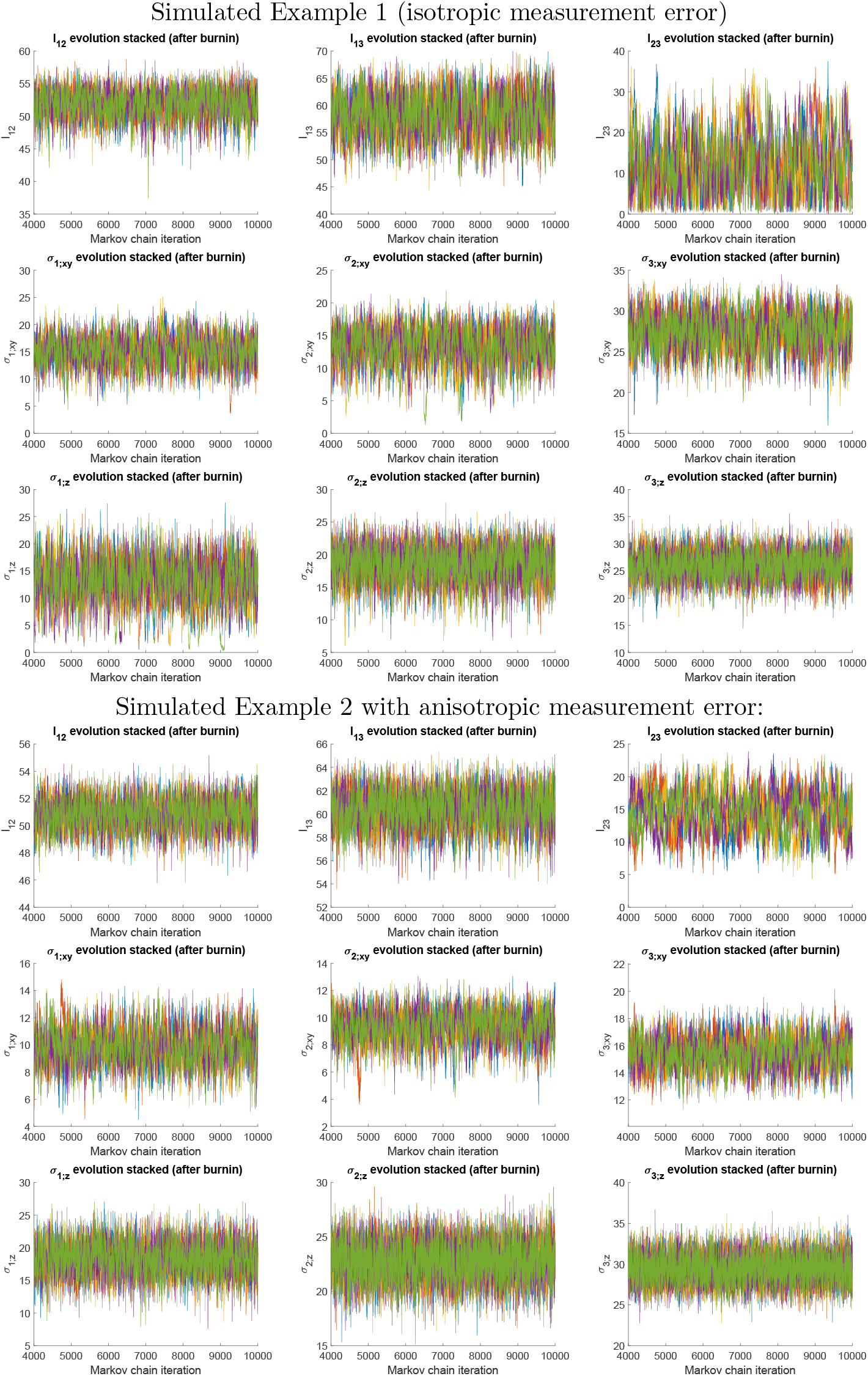
Markov chain traces of each of the parameters *l*_*ij*_ = |*X*_*j*_ − *X*_*i*_|, *σ*_*i*;*d*_ for the single-state simulated Examples 1, 2 in Table 1. Traces are plotted post burnin and sub-sampled to give 10000 samples. Five independent chains are overlain in separate colours.

**Fig S13.2.**
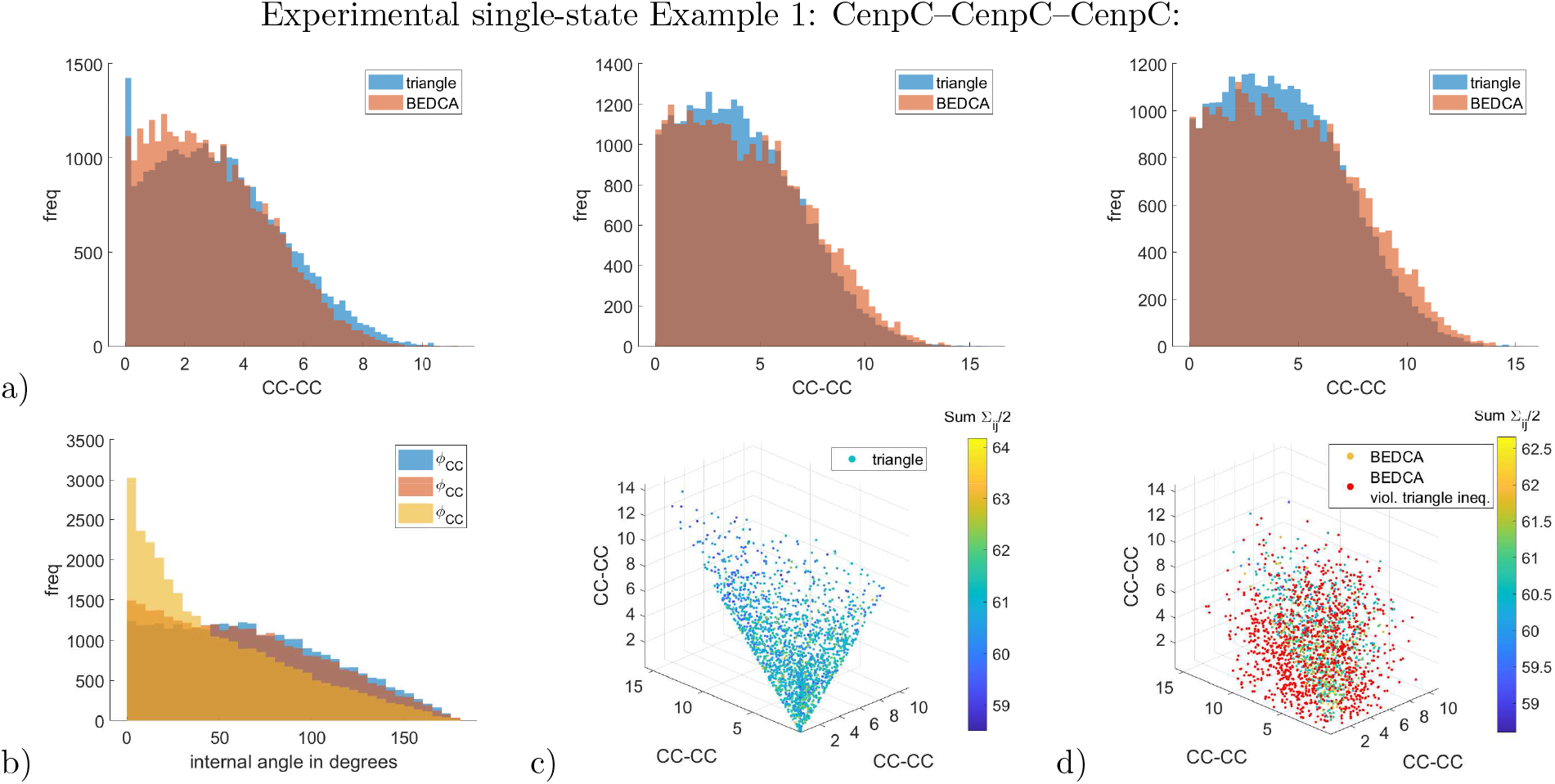
Marginal posteriors of the triple labelled CenpC experiment (Example 1 in Table 3). Panels are the same as in Fig 3. Constructing a joint distribution from the three pair-wisely inferred lengths assuming independence yields 62% violations of the triangle inequality (red dots in panel d)).

**Fig S13.3.**
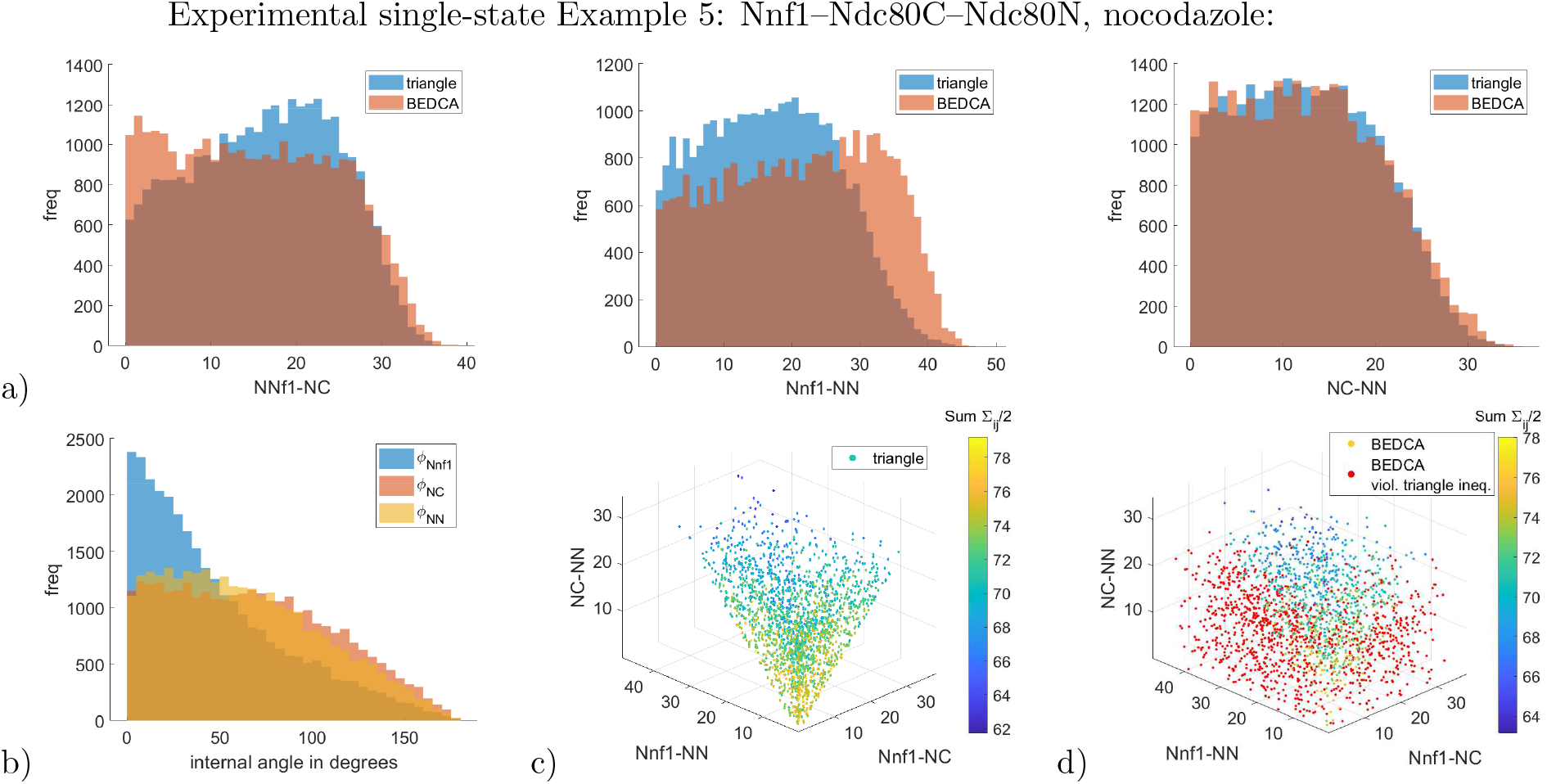
Marginal inferred posteriors of the Nnf1–Ndc80C–Ndc80N experiment in nocodazole treatment (Example 5 in Table 3). Panels are the same as in Fig 3. Constructing a joint distribution from the three pair-wisely inferred lengths assuming independence yields 57% violations of the triangle inequality (red dots in panel d)).

### S14. Additional supplementary tables

**Table S14.1.** Inferred measurement errors *σ*_*j*;*d*_ for single-state simulated Examples 1, 2 in Table 1 for each marker individually. Due to an indistinguishability for the pair-wise methods, these parameters can only be inferred, if at least three markers *J* ≥ 3 are used.

| (lengths and errors in nm) | $\sigma_{1;xy}$ | $\sigma_{1;z}$ | $\sigma_{2;xy}$ | $\sigma_{2;z}$ | $\sigma_{3;xy}$ | $\sigma_{3;z}$ |
| --- | --- | --- | --- | --- | --- | --- |
| Example 1 with isotropic measurement error: |  |  |  |  |  |  |
| true value | 15 | 15 | 15 | 15 | 25 | 25 |
| triangle, uninformative prior | $15.0 \pm 2.7$ | $13.0 \pm 4.0$ | $13.2 \pm 2.8$ | $18.5 \pm 2.5$ | $27.4 \pm 2.1$ | $25.9 \pm 2.4$ |
| Example 2 with anisotropic measurement error: |  |  |  |  |  |  |
| true value | 10 | 20 | 10 | 20 | 15 | 30 |
| triangle, uninformative prior | $9.8 \pm 1.3$ | $18.3 \pm 2.4$ | $9.3 \pm 1.2$ | $22.7 \pm 1.7$ | $15.5 \pm 1.1$ | $29.4 \pm 1.7$ |

**Table S14.2.** Data sizes of experimental examples. The number of kinetochores and number of cells are given after image processing and quality control.

| Single-state experimental examples (see Table 3 for results): |  |  |  |  |  |
| --- | --- | --- | --- | --- | --- |
| Example | triple structure | treatment | number of kinetochores $N$ | number of cells | |
| 1 | CenpC–CenpC–CenpC |  | 1250 | 32 |  |
| 2 | CenpC–Nd80C–Ndc80N | DMSO | 72 | 3 |  |
| 3 | CenpC–Nd80C–Ndc80N | nocodazole | 118 | 5 |  |
| 4 | Nnf1–Nd80C–Ndc80N | DMSO | 570 | 18 |  |
| 5 | Nnf1–Nd80C–Ndc80N | nocodazole | 238 | 8 |  |

| Two-state experimental examples (see Table 4 for results): |  |  |  |  |  |
| --- | --- | --- | --- | --- | --- |
| Example | triple structure | $p_{\text{infused}}^{(2)}$ | number of kinetochores $N_{\text{mix}}$<br>mixture dataset | number of kinetochores $N_{\text{inform}}$<br>informing dataset | number of cells |
| 1 | Nnf1–Ndc80C–Ndc80N | 0.0% | 188 | 238 | 14 |
|  | Nnf1–Ndc80C–Ndc80N | 10.5% | 210 | 216 | 14 |
|  | Nnf1–Ndc80C–Ndc80N | 19.7% | 234 | 192 | 14 |
|  | Nnf1–Ndc80C–Ndc80N | 29.9% | 268 | 158 | 14 |
|  | Nnf1–Ndc80C–Ndc80N | 40.1% | 314 | 112 | 14 |
| 2 | CenpC–Ndc80C–Ndc80N | 0.0% | 72 | 118 | 8 |
| 3 | Nnf1–Ndc80C–Ndc80N | 0.0% | 570 | 238 | 26 |

**Table S14.3.** Inferred measurement errors *σ*_*j*;*d*_ for single-state experimental Examples 1 – 5 in Table 3 for each marker individually. Here we abbreviate CC for CenpC, NC for Ndc80C and NN for Ndc80N. The same fluorophores were used across all examples to mark the various structures of the kinetochore (apart from the triple-CenpC, Example 1, where A568 is the secondary antibody of the CenpC in the other examples). The table shows consistency of the inference results of the same fluorophore between different examples. Here, the marker for CenpC exhibits the smallest measurement error, being up to only half as large as some of the other markers. Slight differences can be observed for the same fluorophores between DMSO- and nocodazole-treated cells (Nnf1, Ndc80N). Due to an indistinguishability for the pair-wise methods, these parameters can only be inferred, if at least three markers *J* ≥ 3 are used.

| (lengths and errors in nm) | $\sigma_{CC;xy}$ | $\sigma_{CC;z}$ | $\sigma_{Nnf1;xy}$ | $\sigma_{Nnf1;z}$ | $\sigma_{NC;xy}$ | $\sigma_{NC;z}$ | $\sigma_{NN;xy}$ | $\sigma_{NN;z}$ |
| --- | --- | --- | --- | --- | --- | --- | --- | --- |
| DMSO-treated experimental single-state examples: |  |  |  |  |  |  |  |  |
| Example 1, A488, triangle, uninf. prior | $13.1 \pm 0.3$ | $25.4 \pm 0.9$ | / | / | / | / | / | / |
| Example 1, A568, triangle, uninf. prior | $9.0 \pm 0.4$ | $20.3 \pm 1.1$ | / | / | / | / | / | / |
| Example 1, A647, triangle, uninf. prior | $16.1 \pm 0.4$ | $40.4 \pm 1.0$ | / | / | / | / | / | / |
| Example 2, triangle, uninf. prior | $9.4 \pm 3.1$ | $16.5 \pm 7.7$ | / | / | $12.1 \pm 1.8$ | $33.6 \pm 4.1$ | $13.0 \pm 2.7$ | $28.2 \pm 6.2$ |
| Example 4, triangle, uninf. prior | / | / | $10.6 \pm 0.8$ | $25.1 \pm 2.0$ | $13.1 \pm 0.6$ | $30.7 \pm 1.5$ | $9.5 \pm 0.9$ | $27.6 \pm 2.0$ |
| Nocodazole-treated experimental single-state examples: |  |  |  |  |  |  |  |  |
| Example 3, triangle, uninf. prior | $13.8 \pm 3.3$ | $16.3 \pm 7.3$ | / | / | $18.2 \pm 2.2$ | $35.5 \pm 3.9$ | $18.4 \pm 2.6$ | $39.1 \pm 4.0$ |
| Example 5, triangle, uninf. prior | / | / | $20.0 \pm 2.7$ | $32.7 \pm 2.7$ | $15.0 \pm 2.5$ | $26.6 \pm 2.7$ | $23.6 \pm 1.8$ | $31.5 \pm 2.6$ |

## Notes

### Competing Interest Statement

The authors have declared no competing interest.

